# A Deep Survival EWAS approach estimating risk profile based on pre-diagnostic DNA methylation: an application to Breast Cancer time to diagnosis

**DOI:** 10.1101/2022.02.25.481911

**Authors:** Michela Carlotta Massi, Lorenzo Dominoni, Francesca Ieva, Giovanni Fiorito

## Abstract

Previous studies for cancer biomarker discovery based on pre-diagnostic blood DNA methylation profiles, either ignore the explicit modeling of the time to diagnosis (TTD) as in a survival analysis setting, or provide inconsistent results. This lack of consistency is likely due to the limitations of standard EWAS approaches, that model the effect of DNAm at CpG sites on TTD independently. In this work, we argue that a global approach to estimate CpG sites effect profile is needed, and we claim that such approach should capture the complex (potentially non-linear) relationships interplaying between sites. To prove our concept, we develop a new Deep Learning-based approach assessing the relevance of individual CpG Islands (i.e., assigning a weight to each site) in determining TTD while modeling their combined effect in a survival analysis scenario. The algorithm combines a tailored sampling procedure with DNAm sites agglomeration, deep non-linear survival modeling and SHapley Additive exPlanations (SHAP) values estimation to aid robustness of the derived effects profile. The proposed approach deal with the common complexities arising from epidemiological studies, such as small sample size, noise, and low signal-to-noise ratio of blood-derived DNAm. We apply our approach to a prospective case-control study on breast cancer nested in the EPIC Italy cohort and we perform weighted gene-set enrichment analyses to demonstrate the biological meaningfulness of the obtained results. We compared the results of Deep Survival EWAS with those of a traditional EWAS approach, demonstrating that our method performs better than the standard approach in identifying biologically relevant pathways.

**Author summary:** Blood-derived DNAm profiles could be exploited as new biomarkers for cancer risk stratification and possibly, early detection. This is of particular interest since blood is a convenient tissue to assay for constitutional methylation and its collection is non-invasive. Exploiting pre-diagnostic blood DNAm data opens the further opportunity to investigate the association of DNAm at baseline on cancer risk, modeling the relationship between sites’ methylation and the Time to Diagnosis. Previous studies mostly provide inconsistent results likely due to the limitations of standard EWAS approaches, that model the effect of DNAm at CpG sites on TTD independently. In this work we argue that an approach to estimate single CpG sites’ effect while modeling their combined effect on the survival outcome is needed, and we claim that such approach should capture the complex (potentially non-linear) relationships interplaying between sites. We prove this concept by developing a novel approach to analyze a prospective case-control study on breast cancer nested in the EPIC Italy cohort. A weighted gene set enrichment analysis confirms that our approach outperforms standard EWAS in identifying biologically meaningful pathways.

## Introduction

DNA methylation (DNAm) is a chemical modification that consists of the addition of a methyl group via a covalent bond to the cytosine ring of DNA, in correspondence of CpG sites (CpGs), and a large body of evidence has demonstrated that CpG islands hypermethylation is implicated in loss of expression of a variety of critical genes in cancer [1].

These alterations can be detected both in the target tissue (e.g., cancer biopsy vs tumor-free adjacent tissue) and in blood-derived DNA. DNAm dysregulation in target tissue are likely the effect of the disease rather than vice versa [2], whereas DNAm alterations in blood are commonly used as biomarkers of long-term exposure and insults to the DNA which includes variability related to genetic predisposition or individual response to risk factors. The above suggest the possibility to use blood-derived DNAm profiles as new biomarkers for cancer risk stratification and possibly, early detection. Investigating DNA methylation data from blood samples is of particular interest since it is a convenient tissue to assay for constitutional methylation and its collection is non-invasive. Moreover, exploiting pre-diagnostic blood DNAm data opens the opportunity to investigate the association of DNAm at baseline on cancer risk, modeling the relationship between sites’ methylation and the Time to Diagnosis (TTD). That would indeed be desirable, as to improve the effectiveness of current screening procedures via the definition of novel effective and non-invasive biomarkers (e.g., via DNAm-based scoring methods) is a public health necessity.

In this work, we focus on the identification of blood DNAm profiles predictive of TTD, with the aim to improve the reliability/reproducibility of the results, as well as their biological meaningfulness. Indeed, previous studies based on pre-diagnostic blood DNAm, either ignore the explicit modeling of TTD as in a survival analysis setting, or mostly provide inconsistent results (e.g. [3] and references therein). This unsatisfactory outcome may be induced by the limitations of the most traditional approaches for the analysis of whole-genome DNAm data, i.e., Epigenome-Wide Association Studies (EWAS). Indeed, EWAS analyses traditionally comprise multiple independent tests of individual CpG sites or regions, seeking for significant associations by imposing p-value thresholds corrected for multiple comparisons.

The main limitations of this approach are: (i) the extremely high dimensionality of epigenome-wide DNAm data affects the reliability of multiple testing correction, driving p-value thresholds down to extremely low values, (ii) the strong correlation among methylated sites, that is usually not considered in statistical modelling, (iii) the presence of several (likely unmeasured) confounders, since DNAm profiles are influenced by environmental exposures and lifestyle behaviors [4]. Additionally, the context of pre-diagnostic blood DNAm carries further complexities due to the (iv) very low signal-to-noise ratio of differential methylation, as both cases and controls are healthy at the time of DNAm collection. Lastly, (v) these methods based on independent testing do not account for any of the complex and potentially non-linear interactions that might exist between CpG sites or the combined effect of multiple loci together on the phenotype. Indeed, findings from previous studies [5], suggest the need for an epigenome-wide approach, that assigns individual parameters while accounting for sites’ combined effect on the phenotype. We refer to this set of parameters as an effects profile. Nonetheless, exploiting biostatistical approaches s.a. Cox Proportional Hazard (CoxPH) regression, to model survival outcomes and infer this effect profile including all CpG sites as predictors together, would lead to further methodological pitfalls. Firstly, modeling such a large number of covariates leads to effect size overestimation. Moreover, CoxPH models suffer the multi-collinear nature of CpG sites and are based on strong assumptions, such as the additive nature of covariates’ effect on the outcome, unless including an even larger number of terms to account for interactions. These limitations and the complexities of DNAm data can be naturally handled by Machine Learning (ML) approaches, such as Neural Networks (NN) and Deep Learning (DL)-based methods. Indeed, NN are optimized to extract rich latent features from DNAm data, handling multi-collinearity, noise, and considering the complex non-linear interactions between very large amounts of input covariates [6]. Some recent works demonstrated the usefulness of AutoEncoders (AE), Variational AE (VAE) and DL models to obtain DL-based EWAS (henceforth, Deep EWAS) [7–9], especially when coupled with post-hoc Explanation Methods (EM) [6]. EM like SHapley Additive exPlanations (SHAP) [10,11] try to overcome the “black box” aspect of these complex models assigning a contribution score (i.e., an importance weight) to each input feature, based on how much it contributed to the model prediction. In Deep EWAS, EMs are exploited to provide insightful information on how DNAm input determine the outcome, and the weights can be considered as an estimation of the aforesaid effects profile on the phenotype. Nonetheless, the highly parametrized deep models are prone to overfitting, unless they are presented with very large training samples, and a suboptimal training may result in unstable and unreliable explanations (i.e., importance weights). Indeed, to obtain effective explanations to derive meaningful conclusions from, both model’s accuracy and importance weights stability should be maximized [12].

While ML and DL approaches to EWAS are gaining momentum, the task of modeling TTD with the objective of inferring robust Epigenome-Wide DNAm effects profiles in a survival setting has been largely unexplored and there has not been any application yet in the context of blood DNAm. Indeed, to the best of our knowledge, the existing Deep EWAS literature mainly focuses on classification settings (e.g., cancer type, cancer status, patients’ clinical condition, etc.). The most prominent examples of this effort come from the works from Levy et al. [6], that, in its seminal work, proposes an effective Deep EWAS framework based on VAE-based encoding of DNAm data, followed by a prediction model explained through SHAP. Despite the flexibility of the algorithm, no effort has been devoted to tackle the specific facets of a time to event setting. Only one related study [8] deals with DNAm data and time of BC recurrence to filter significant latent features. Yousefi et al. [13] propose a general framework for genome-wide data, but their automatic hyperparameter optimization does not account for the stability of their back-propagation based explanations. Despite the lack of efforts in modeling blood-derived DNAm via modern ML-based techniques, we argue that the search for effective prognostic biomarkers from this type of data would instead benefit from a paradigm shift from standard EWAS approaches. Indeed, we believe that to achieve reliable and biologically meaningful results in the search for TTD biomarkers from blood-derived DNAm data, a global approach to estimate CpG sites effects profile is required. Furthermore, we claim that such approach should exploit the potential of DL-based methods to capture the complex and potentially non-linear relationships interplaying between sites, after a proper tailoring of the model to deal with time-to-event outcomes and the facets of real-life epidemiological studies.

In this work, we validate our hypothesis by analyzing blood based DNAm data from a prospective case-control study on Breast Cancer (BC) nested in the EPIC Italy cohort. This cohort, that was previously analyzed in [3,5], presents all typical complexities of pre-diagnostic DNAm studies, i.e., small sample size, risk of confounders effect due to the long time from recruitment to cancer diagnosis, and the challenge of identifying differentially methylated sites among individuals that are healthy at the time of blood collection.

To tackle this data and prove our concept, we develop a new DL-based approach that assesses the relevance of individual CpG Islands in determining TTD while modeling them all together in a survival analysis setting. The methodology is inspired by previous Deep EWAS approaches [6,8,14,15] and adapted for survival data, therefore we name it Deep Survival EWAS. Our Deep Survival EWAS models the complex relationships among CpG sites and between sites and TTD, and ultimately estimates the desired CpG Islands effects profile through SHAP, in terms of importance weights in determining the hazard rate.

In summary, the objectives and contributions of this study are multiple:

- We highlight the need of novel approaches to cancer TTD modeling from blood-derived DNAm. To derive meaningful and robust biological insights the proposed method overcomes the limitations of standard EWAS approaches and take into account the combined effect of DNAm sites on cancer onset.
- We present our original approach to the problem, i.e. Deep Survival EWAS. Besides its natural capability to model complex and potentially non-linear interactions among CpG Islands, the overall algorithm has several valuable methodological details meant to deal with the complexities arising from this crucial real-life biological research context, s.a. small samples, noise, and low signal-to-noise ratio of blood-derived DNAm.
- We validate the aforementioned hypothesis presenting the results of our Deep Survival EWAS in an in-depth analysis of a prospective case-control BC study nested in the EPIC Italy cohort. To demonstrate the biological meaningfulness of the obtained effect profile, we perform weighted gene-set enrichment analyses (GSEA), comparing the results of Deep Survival EWAS with those of a traditional EWAS approach. The GSEA results, indicate that our method performs better than the standard approaches both looking at the biological reliability and the statistical stability of genes and pathways identified.
- Finally, we confirm the value of a DL-based model accounting for predictors interactions comparing our method with a simpler Cox model with additive effects only.

## Results

### DNAm data preprocessing

In this study we exploit blood based DNAm data collected for a BC study nested in the EPIC Italy cohort. After data preprocessing, sample and probe filtering our primary dataset includes DNAm values for 13,499 CpG sites in 248 incident BC cases and one-to-one matched controls. DNAm values are expressed as the ratio of methylated cytosines over total cytosines (named from here on *β* values). DNAm *β* values for CpGs pertaining to the same CpG island (according to the UCSC annotation [16]) were grouped as described in Methods section, where we refer the reader to for an in-depth description of the data preparation pipeline (cf. EPIC Breast Cancer Data). The final dataset is composed of DNAm data for 3,807 CpG islands.

### Overview of the Deep Survival EWAS approach

To enhance the extraction of relevant information from blood DNAm data, we defined our methodological approach, i.e. the Deep Survival EWAS algorithm, as a set of steps tailored to face the aforementioned facets of these complex dataset and research settings. In Fig 1, we represent the visual schema of the overall procedure.

**Fig 1.**
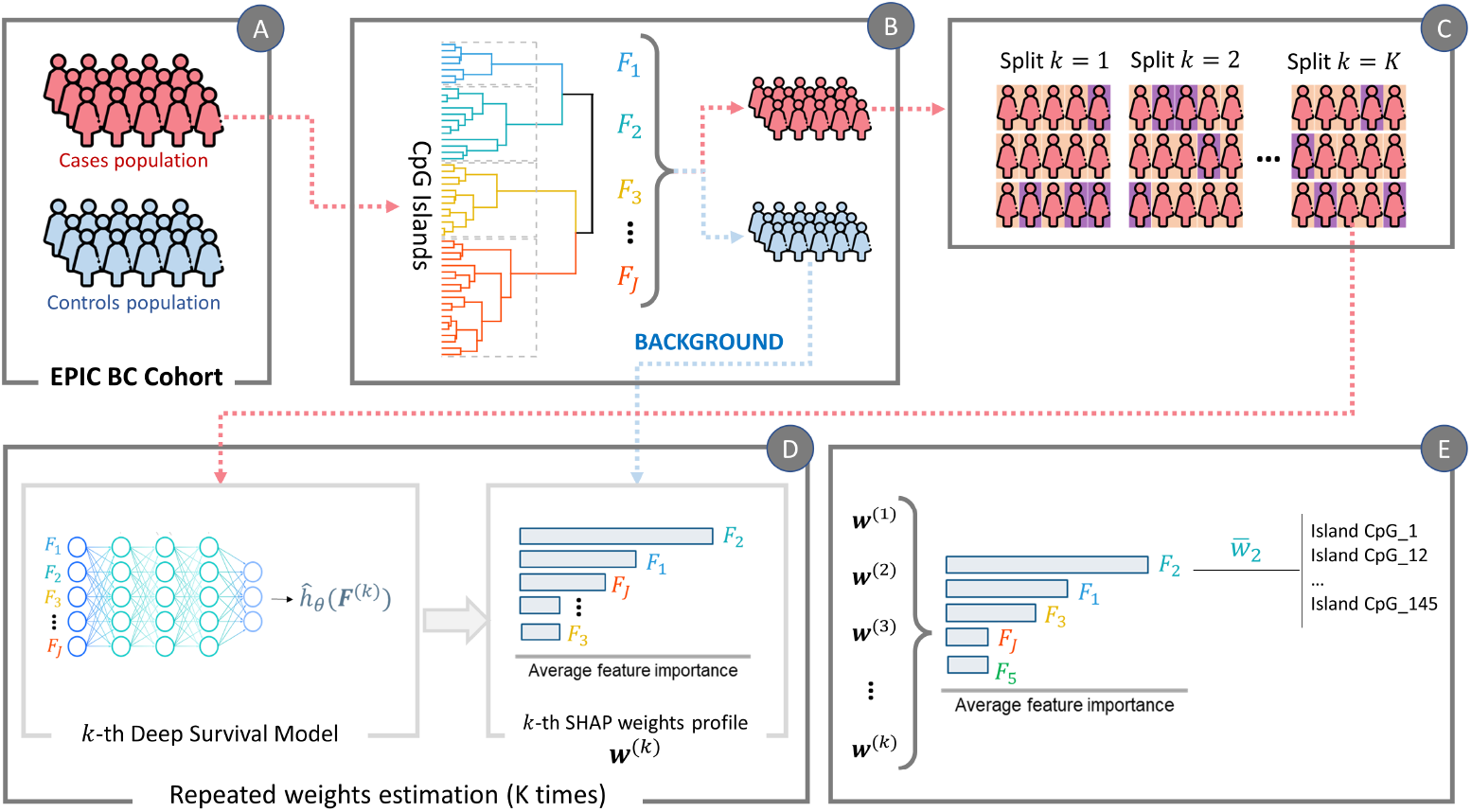
Algorithm pipeline of the methodology applied in this study. **(A)** We started from the EPIC BC cohort, equally split between cases (i.e., patients enrolled healthy but diagnosed with BC within the study time frame) and matched controls (i.e. patients matching cases at baseline, that were not diagnosed with BC within the study time frame). **(B)** The first step is feature aggregation via hierarchical clustering, exploiting CpG Islands continuous *β* values of the population of cases to infer the J clusters of features. The same clustering structure is then applied to both cases and controls, grouping their CpG Islands accordingly. **(C)** The cases population is exploited to generate K independent and randomly split training and test sets, each of them with 80% patients in training set and 20% patients in the test set. **(D)** Each of the K splits goes through step D independently. In particular, the k-th training set is used to train a Deep Survival Model, that then is used to estimate SHAP weights profiles (*w*^*k*^) on the k-th test set using the controls’ population as background data. **(E)** After generating independently K sets of weights profiles, they are aggregated to obtain the final estimation of the effects profile for BC TTD.

In brief, as a first step, we subdivide the study population between cases (i.e., individuals diagnosed with cancer within the study follow-up) and controls (i.e., individual remained healthy until the end of follow-up). We begin with exploiting the population of cases, and we agglomerate CpG Islands through Hierarchical Feature Ranking based on Euclidean Distance of *β*-values. Once the clusters are defined, we aggregate cluster-specific *β*-values computing their mean. Through this step, we obtain a feature matrix *F*_*cases*_ ∈ ℝ^*N ×J*^, where *N* is the number of cases and *J* is the number of feature clusters. Henceforth, these aggregated CpG Islands will be referred to as *features F*_*j*_. We apply the same clustering structure to the CpG Islands of the controls’ population to obtain *F*_*controls*_ ∈ ℝ^*M ×J*^, where *M* is the number of controls.

Then, to make a robust and reliable estimation of the EWAS weights associated with the input *F*_*cases*_, we generate *K* random splits of the dataset into training and test set, with 80/20 ratio and for each training-test split: (i) we model the complex non-linear relationship between the input and TTD, by exploiting a deep survival feed-forward neural network predictive of the log-risk function; (ii) we exploit Kernel Shap from SHAP framework [10] to estimate weights 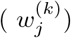 associated to each feature *f* ∈ *F*. Notably, SHAP relies on the use of a “background dataset” to estimate the expected model output and estimates individual feature’s impact on prediction as their contribution to the difference between the observed and the expected prediction (more details in the following section). Therefore, choosing the right background data is crucial to obtain contextually meaningful explanations. Indeed, to resemble the differential estimation of DNAm effects in EWAS, we compute the importance of the features in *F*_*cases*_ using *F*_*controls*_ as reference for SHAP.

### Deep Survival EWAS on the case-control study of Breast Cancer nested in the EPIC Dataset

We applied the just described Deep Survival EWAS to the BC sample from EPIC Italy, to infer an effect profile associated to the blood-derived DNAm CpG Islands in the dataset. As many ML or DL based algorithms, our approach comprises several building blocks that require specific choices in terms of hyperparameters and/or implementation details, that need to be optimized to provide a robust estimation of the desired effect profiles. Indeed, the better the underlying K models will perform in predicting the survival outcome, the more meaningful the feature importance weights derived by SHAP will be. Concurrently, a stable feature importance ranking suggests that the K models trained on different data subsets are consistently capturing and exploiting the information from a specific set of features to obtain such prediction. A satisfactory performance on both aspects together translates in a trustworthy and effective effects profile estimation [17]. Considering the above, we first needed to select the optimal number of feature clusters (J) and identify the best Deep Survival model in terms of architecture and hyperparameters. These two aspects were jointly optimized to maximize the time to event prediction on the population of cases, averaged across *K* = 10 random splits, and the robustness of the K derived effects profile. The performance for the time to event prediction was measured with the Harrel’s Concordance Index (CI), while the robustness of profiles with the Kendall Tau Ranking Stability (KT-stability). Further details on the best model selection and performance metrics’ definitions, including rationale for our choices, can be found in the Methods Section, whereas complete results are available in Supporting Information, S1 Table. The final results presented here, corresponding to the chosen best Deep Survival EWAS configuration, were obtained by grouping CpG Islands into J=128 clusters. The latter were used as the input for the Deep Survival NN model with a J-dimensional input layer followed by three fully connected layers of 64, 32, and 16 nodes respectively. This model resulted in an average CI of 0.702 ± 0.019 and an average KT-Stability of 0.669 ± 0.036.

In Fig 2, left panel, we show the features that correspond to the top 10 highest values in the obtained effects profile on log risk prediction. As mentioned in the previous section, these weights 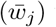 were derived as the average impact on log risk prediction of each feature across the K = 10 random resamples. As for the background dataset, in this case impact was measured w.r.t. the control group of all BC controls. We performed some post-hoc analyses to showcase the tight relationship between the most relevant features according to the estimated effects profile and TTD. In Fig 3, panel A, we split the population of cases into time-to-event classes (i.e., early event [0 − 3.5*y*], mid-early event [3.5*y* − 7*y*], mid-late event [7*y* − 10.5*y*] and late event [10.5 − 16*y*]) and we plot the distribution of the aggregated *β* values of the most important feature (*F*_120_). The boxplot shows a decreasing trend of the feature value with increasing TTD. Note that *F*_120_ groups 20 CpG Islands (cf. Table 1), meaning that higher methylation values on those 20 sites are associated with earlier diagnosis (i.e., higher log risk).

**Fig 2.**
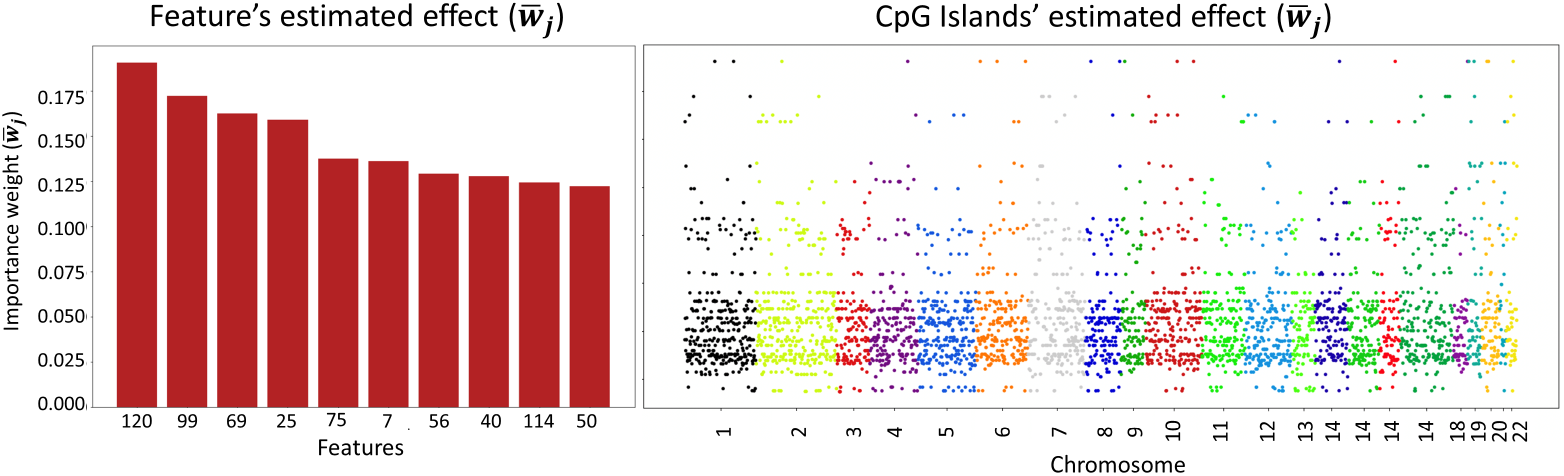
Deep Survival EWAS estimated effects profile. On the left panel, the top 10 aggregated features with the highest associated 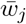 value in the effects profile at Feature’s level. On the right, sharing the same y-axis, the effects profile at CpG Island, where each methylated islands in the dataset is associated with the 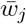 value of the feature they are clustered in.

**Fig 3.**
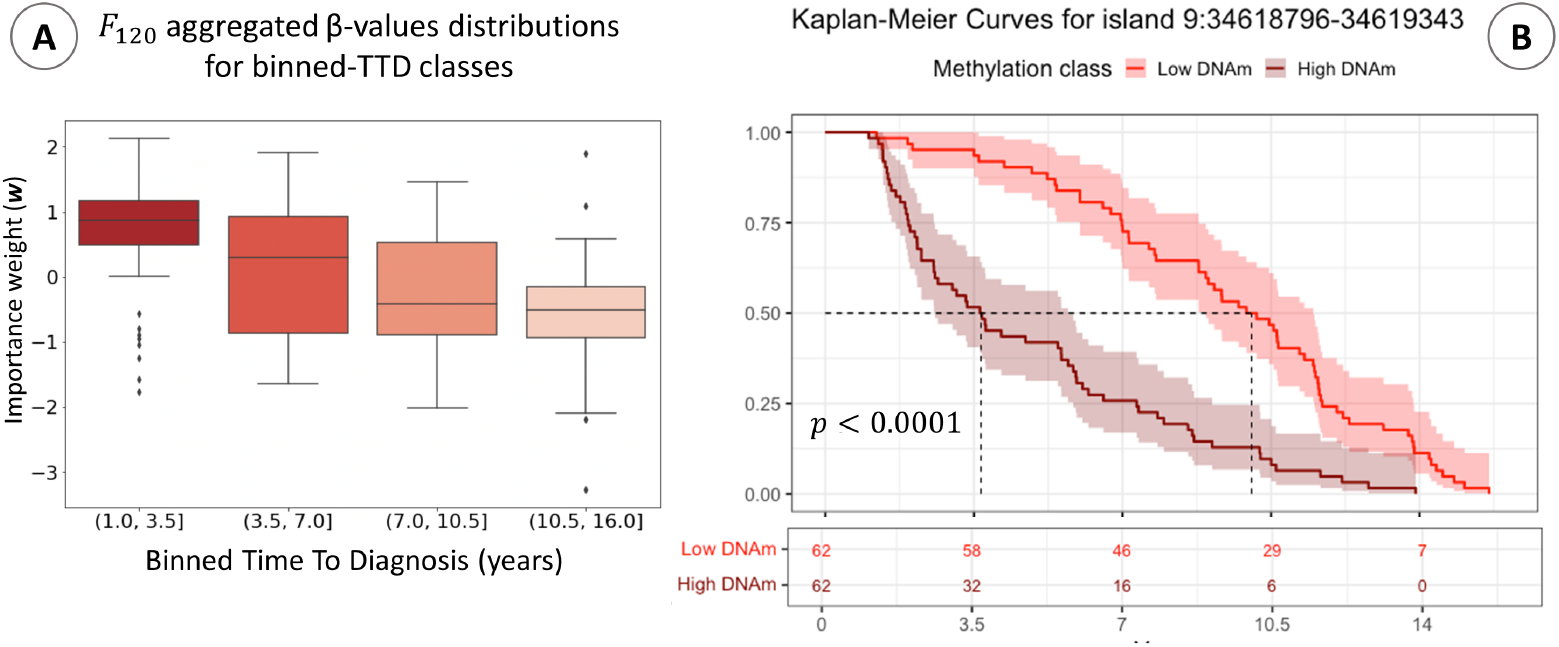
Post-hoc analysis results. **(A)** Distribution of aggregated DNAm beta-values in the feature (*F*_120_) associated with the highest impact in the estimated effect profile. Subjects are binned according to their TTD into four classes. **(B)** Kaplan-Meier curves of low DNAm (below 25th percentile of *β*-values distribution) and High DNAm (above 75th percentile) populations’ in CpG Island 9:34518796-34619343. This island belongs to the cluster that is aggregated into feature *F*_120_. The plot reports the Log-Rank test p-value for the difference between the two groups; the lower part of the plot reports the count of subjects in High DNAm and Low DNAm populations for CpG Island 9:34518796-34619343, according to TTD.

**Table 1.** CpG Islands in Feature 120. List of CpG Islands agglomerated in *F*_120_, therefore assigned to the highest effect weight.

| CpG Island |
| --- |
| 1:90945518-90945656 |
| 1:158090642-158091676 |
| 10:102493904-102494072 |
| 10:119294070-119294143 |
| 14:87862626-87863008 |
| 16:85096322-85097146 |
| 18:75811758-75814395 |
| 19:1704275-1706659 |
| 19:13070446-13070515 |
| 2:100086548-100088317 |
| 20:21438169-21438255 |
| 20:21449303-21449404 |
| 22:37180713-37182260 |
| 4:149584089-149584799 |
| 6:1570179-1570756 |
| 6:43530362-43531683 |
| 6:166137998-166138866 |
| 8:21701267-21701566 |
| 8:145119282-145120028 |
| 9:34618796-34619343 |

As a further validation, for each CpG Island pertaining to *F*_120_, we extracted two sub-population of cases: the hypermethylated population, composed of patients with *β* value above the 75^*th*^ percentile for that CpG island, and the hypomethylated population, composed of patients with values below the 25th percentile. Then, we compared the risk profiles of the two groups via Kaplan-Meyer (KM) test. In Fig 3, panel B, we plot the KM curve obtained and the log-rank test p-value for CpG Island 9:34518796-34619343, as representative of the 20 islands in F120. Similar behavior and log-rank test results was obtained for the other 19 islands in F120 (all KM curve plots for the CpG Islands in *F*_120_ are reported in S1 File). As expected, modeling TTD of BC cases only, both KM tend to 0 as t increases. However, it is clear from KM and test results how methylation on those sites is associated with TTD for those patients. It is crucial to note that, despite modeling cases TTD only, the inference of features’ relevance exploits data form the controls group used as the reference (i.e. background). In other words, CpG Islands associated with higher weights by our Deep Survival EWAS approach are those with higher influence on the difference of risk prediction (hazard rate) compared to an expected model built with methylation profiles from the control group. Finally, the right panel of Fig 2 represents the resulting effects profile in terms of CpG Islands, that is the final goal of our Deep Survival EWAS. Specifically, to this end we associated each site to the weight of the feature it was clustered in. The whole list of the ranked features *F*, with the respective CpG Islands they cluster and the resulting effects profile can be found in S2 Table (first sheet for BC controls’ reference group).

### Validation through Gene Set Enrichment Analysis

To validate the biological relevance of the identified effects profile, we performed a Gene Set Enrichment Analyses (GSEA) using a Weighted Kolmogorov-Smirnov (WKS) test [18] (cf. Methods). For each CpG site, the weight (input for the enrichment analysis) is that of the feature *F*_*j*_ the CpG site belongs to. In Fig. 4 we present the KEGG pathways enriched according to the WKS procedure. We found 12 pathways with empirical p-value (1,000 permutations) lower than 0.05, being ‘Pathways in Cancer’ the most significant (empirical *p <* 0.0001). Interestingly, all the identified pathways were previously described as dysregulated in breast cancer, including ‘Human Papilloma virus infection’ [19] and ‘Epstein-bar virus infection’ [20] in addition to well-known BC related pathway like ‘Breast Cancer’, ‘PI3K/Akt/mTOR signalling pathway’, ‘Calcium Signaling pathway’ and ‘Mineral absorption’ [21,22], and some cancer generic pathways like ‘Signaling pathways regulating pluripotency of stem cells’, ‘Proteoglycans in cancer’, ‘Neuroactive ligand-receptor interaction’, and ‘Cytokine-cytokine receptor interaction’. As mentioned, the inference of feature importance, and the derived effects profile, can be influenced by the chosen *background*. Thus, different biological insights can be gathered by tailoring the reference group to answer different research question. For this study we wished to investigate both the robustness of the enrichment results changing the background sample, and whether some additional biological associations could be collected on EPIC BC case-control study estimating weights 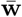 according to different *background* control groups. Therefore, we created 3 additional reference groups to use as background:

- **’BC Matching Controls’** was defined specifically for each of the K splits. In other words, for the k-th test set including 20% of BC cases, we defined the k-th BC Matching Controls group including only the matched controls of those cases. Therefore, we had K BC Matching Controls datasets (with potential subjects’ overlap) to use as background data in estimating effect profiles.
- **’All Controls’** sample included all female control subjects collected for breast, lung and colon cancer. It contained 556 healthy individuals supplied together as Background data.
- **’All Controls with Cases’** included all subjects of the previous sample, with the addition of female cases diagnosed with lung or colon cancer during the EPIC follow up period.

**Fig 4.**
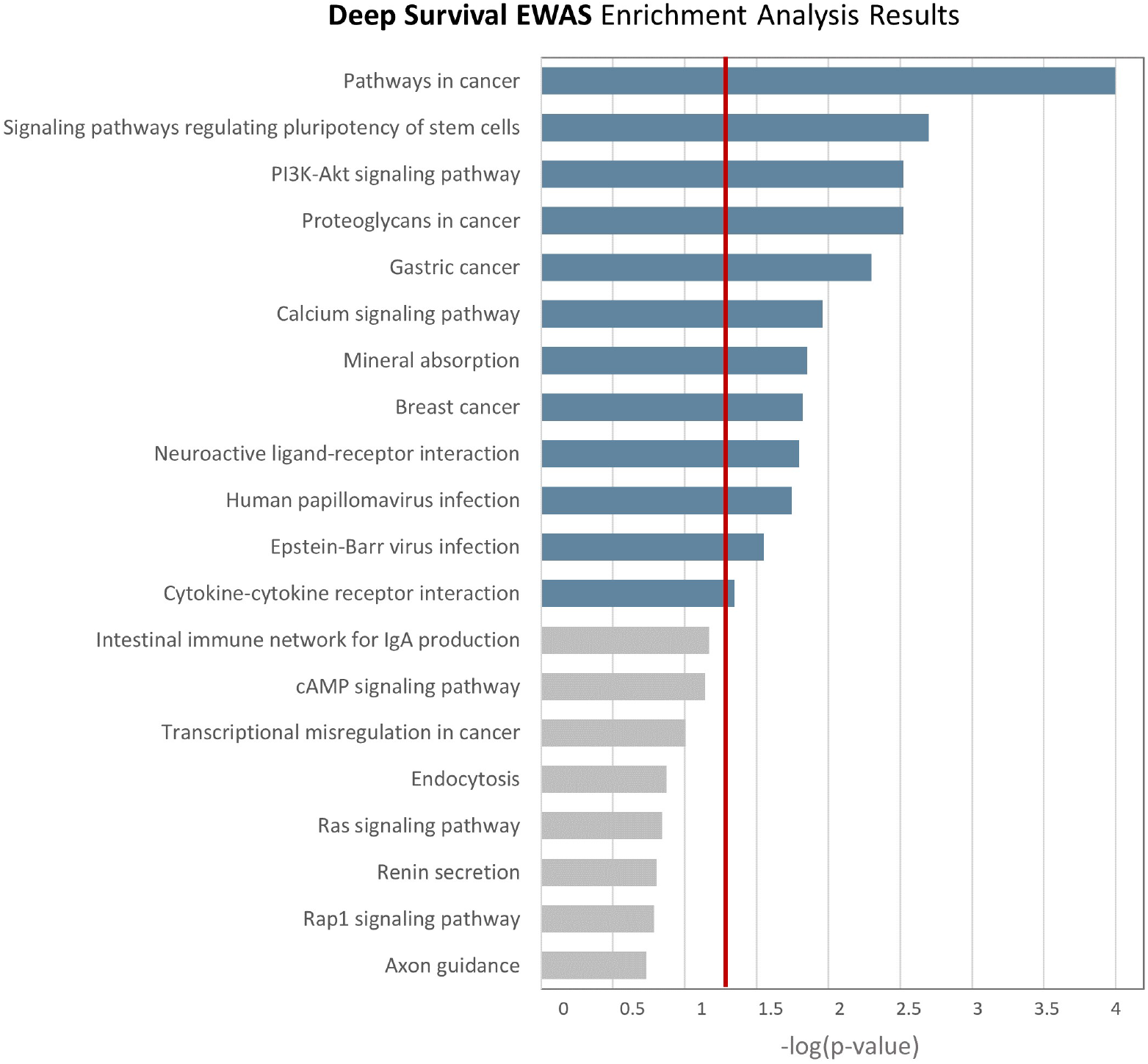
Enrichment analysis results of Deep Survival EWAS on Breast Cancer Controls reference group. In blue are highlighted the significantly associated pathways, i.e. those with empirical p-value above 0.05 (red vertical line), estimated through 10,000 permutations.

To perform these additional analyses, we kept the same optimal configuration of Deep Survival EWAS, supplying to SHAP applied to the survival NN different background samples to estimate *w*_*j*_ for the 128 features. In Fig. 5, we report the results of the three additional enrichment analyses. The full tables of enrichment analyses results for all four reference groups can be found in Supporting Information S3 Table.

**Fig 5.**
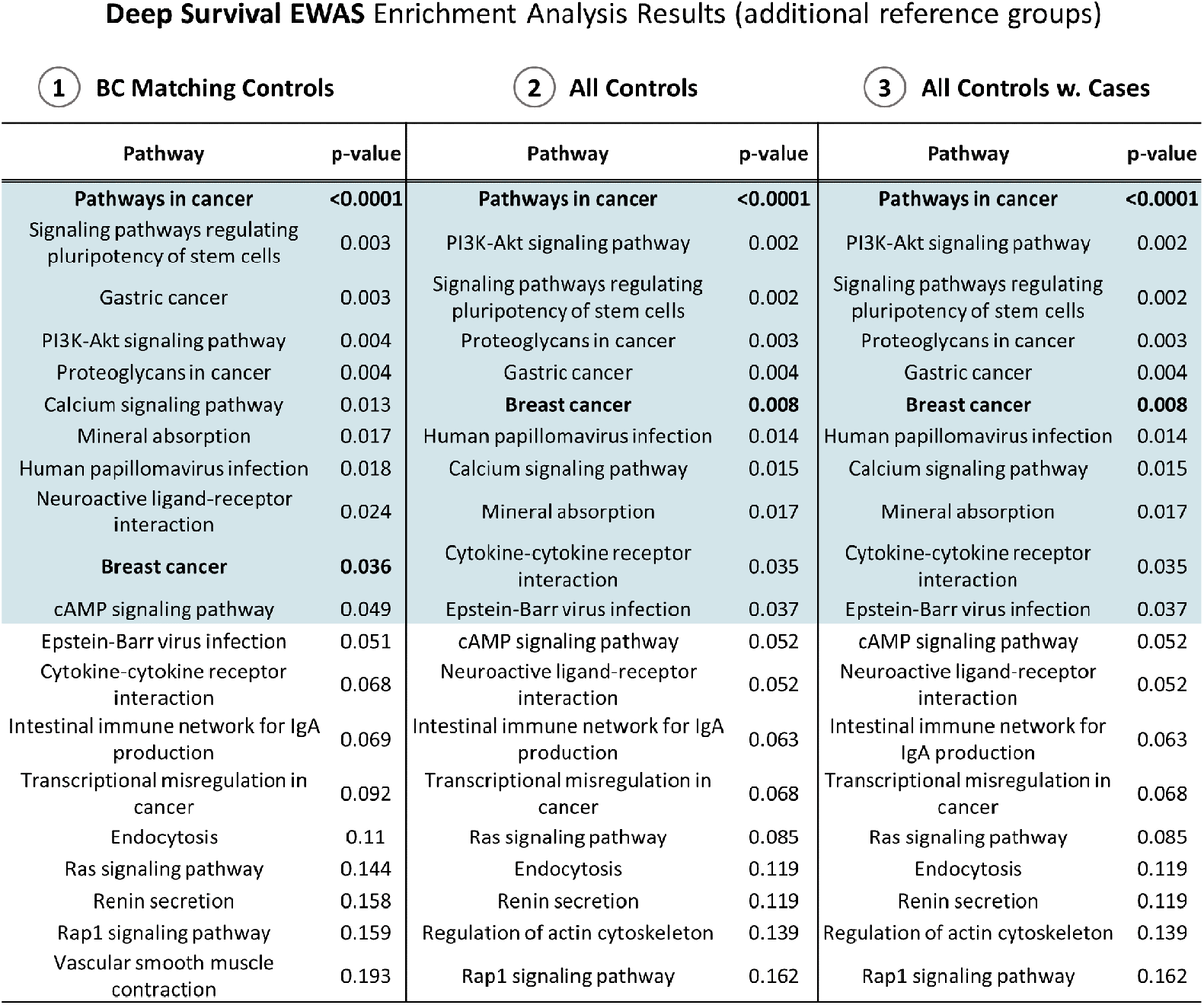
Weighted enrichment analysis results from Deep Survival EWAS (additional reference groups). In blue are highlighted the pathways with significant association, i.e. empirical p-value based on 10,000 permutations.

### Deep Survival outperforms Standard EWAS in identifying biologically meaningful effects profiles

In this work we argue that a complex DL-based approach to estimate a global effects profile on TTD from blood-based DNAm can provide better biological insights compared to traditional approaches. Therefore, to prove our concept and investigate whether taking into account the complex interrelationships among features and outcome modeled with Deep Survival leads to more relevant discoveries than Standard EWAS, we compared the enrichment analyses of the two approaches. In Fig. 6, we report the results of the weighted GSEA for Standard EWAS, where weights were estimated independently for each CpG Island as the test statistic of a univariate Cox Survival Model. For consistency with our approach, we modeled each univariate CoxPH for the cases population only. Detailed tabular results (i.e. p-values and test statistics) are reported in Supporting Information S4 Table). The deriving estimated weights profile (i.e. all CpG Islands p-values, with or without Bonferroni adjustment) are represented in S1 Fig. Finally, Standard EWAS GSEA tabular results are reported in S5 Table. We found eight pathways with empirical p-value lower than 0.05, being ‘Non-small cell lung cancer’ the most significant. Among the identified pathways, two of them are related with the immune system regulation ‘Intestinal immune network for IgA production’ and ‘Chemokine signalling pathway’; two of them with inflammatory processes ‘Leukocyte transendothelial migration’ and ‘C-type lectin receptor signalling pathway’, whereas none of them have been previously described as BC specific.

**Fig 6.**
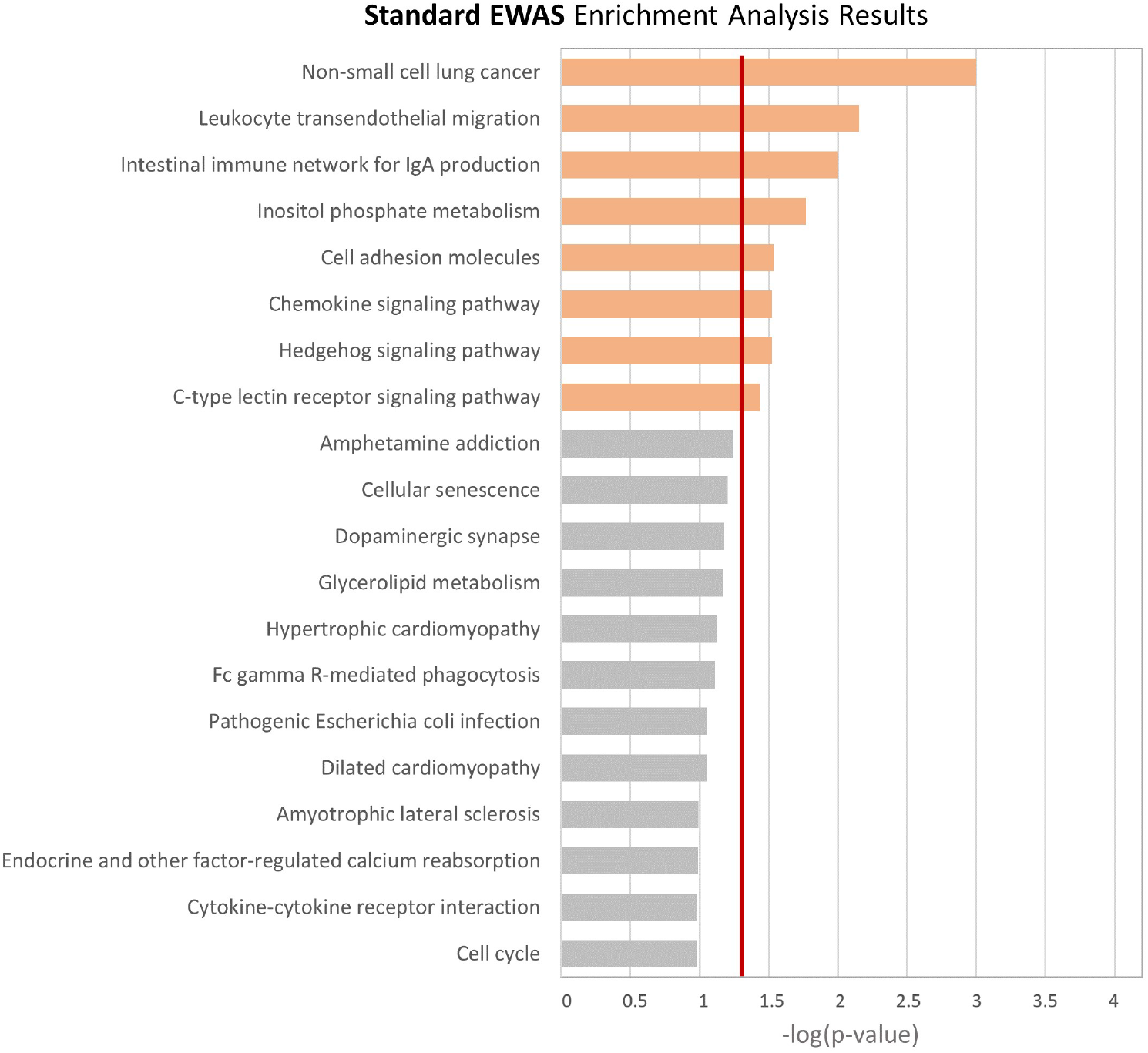
Weighted enrichment analysis results from Standard EWAS approach

### The value of complex non-linear modeling of CpG sites interactions

So far, we validated our claim on the need for a global approach to blood-based DNAm. In our Deep Survival EWAS, this global view is obtained by modeling complex and potentially non-linear interactions between DNAm sites via a deep non-linear survival NN. Nevertheless, a global approach can, by definition, be estimated with a simpler model accounting for the effects of all CpG Islands together with additive contribution on the phenotype only. An example of such a simpler model is a CoxPH model for TTD with a multivariate input and no interaction terms. Therefore, to justify our choice of including a more complex Survival NN model in our procedure, we tested a multivariate epigenome-wide CoxPH model against the two metrics we deem relevant to evaluate the reliability of the derived effects [12]: model prediction performance and robustness of the estimated effects across the *K* subsamples, using the C-index and KT stability metrics respectively. For consistency, we supplied as input to the CoxPH model the CpG Islands aggregated in the same features exploited by our Deep Survival EWAS approach. As detailed in the Methods Section, we tested the performance of the CoxPH model for all feature aggregation (*J*) tested when optimizing our approach. To compute CoxPH KT-stability, the weights rank was derived from the regression parameters (i.e., the effect size) of the CoxPH model. Complete results are reported in S6 Table in Supporting Information. For all values of *J*, our approach outperforms CoxPH on both metrics, especially for the CI (cf. S2 Fig and S3 Fig).

## Discussion

In this work, we presented our approach to solve the complex task of determining the effect of pre-diagnostic blood DNAm in prospective epidemiological studies. Specifically, we focused on the time from recruitment to cancer diagnosis outcome, with the aim of modeling the effect of DNAm CpG sites on the phenotype as in a survival (time to event) analysis setting. To demonstrate the need of a paradigm shift from Standard EWAS approaches (i.e. multiple independent tests) for the analysis of blood based DNAm, we developed and applied a novel approach, Deep Survival EWAS, that robustly estimates weights associated to each CpG Island by considering the combined (global) effect of CpG islands and their complex interactions on the phenotype. To this aim, we modeled the non-linear relationships and interplay between CpG sites and TTD by first grouping them into aggregated features and then feeding them as input to a non-linear deep survival NN, deriving the effects profile through SHAP. We validated our claims analyzing pre-diagnostic blood-derived DNAm from a BC case-control study nested in the EPIC Italy cohort. This is the first attempt to analyze a dataset from an epidemiological prospective study trying to model explicitly the TTD using a DL approach. Notably, BC research is one of the areas that could mostly benefit from a proper modeling of TTD from blood based DNAm, as well as one that suffered the most from inconclusive outcomes (e.g. [3] and references therein). This makes the case study presented in this work an interesting yet challenging testing ground for our proof of concept.

By estimating the desired global effects profile for BC TTD on EPIC DNAm data, we note that the CpG islands grouped in the most relevant features show a clear association of blood DNA methylation with the TTD, with higher methylation values associated with a higher risk of BC in the short term, as shown in the KM curves. Moreover, the overall biological meaningfulness of our procedure was confirmed by the results of a GSEA that identified pathways previously described as associated with cancer onset (with some BC-specific pathway). Instead, a GSEA analysis based on the results of a Standard EWAS (one association test for each CpG island) identified molecular pathways indirectly associated with cancer onset (via immune system dysregulation or inflammatory processes), whereas none of them was BC specific. This results suggest how a global model of methylation profile captures the relationship with the phenotype better than considering DNAm variables one-by-one.

Then, we compared the survival modeling and weights’ reliability performances (i.e., CI and KT-stability) of our approach against a multidimensional CoxPH model, demonstrating that even after CpG islands aggregation into features clusters, a global yet simpler additive effects-based model could not obtain comparable results, further supporting the need to account for non-linear interactions among DNAm predictors as well.

Besides the applied methodology is computationally more expensive than Standard EWAS, or more parametrized and complex than a more traditional survival model, the above results provide strong evidence about the advantages of using such an approach in Epigenome-Wide prospective studies using blood DNAm data. Additionally, we compared the results from GSEA by varying the reference group. This is the first time this peculiarity of SHAP is exploited to gain more insights from a biological perspective, comparing the biological function of the different weights associated to CpG Islands. Specifically, we did not observe significant differences when including women who developed other type of cancers (lung and colon) within the control group. These results suggest our findings provide a specific DNAm signature of BC cancer risk rather than a DNAm signature for all cancer risk.

Whilst we believe the most relevant highlight of this study lies in Deep Survival EWAS achievement of an improved biological interpretability of blood-derived DNAm data on TTD, the tailored choices we made in algorithm design deserve some attention as well. First of all, we decided to focus on modeling TTD for BC cases only. This approach resembles the case-only analysis performed through a standard EWAS approach in [23]. From a methodological standpoint, it allowed to improve survival modeling accuracy, increasing the reliability of the derived explanations [17]. Other methodological details were included to account for all the complexities and peculiarities of both DNAm data facets, and real-life research settings. In particular, clustering CpG Islands based on similarity had the objective of reducing noise and dimensionality simultaneously. The latter aspect reduces the effort in parametrizing the downstream survival model based on NN, alleviating the risk of overfitting on very small sample size, and obtaining suboptimal training results. Besides, this step had a biological justification in that previous studies identified potentially non-contiguous genetically controlled methylation clusters significantly associated to several diseases [24]. Furthermore, the rationale for the metrics exploited in CpG Islands clustering (i.e. Euclidean Distance and mean) was inspired by the results presented in Gagliardi et al. [5], where the association between blood-based DNAm and the phenotype was modeled under the hypothesis that extreme methylation values (i.e., epimutations) are significantly associated with the outcome. The Euclidean Distance would first identify Islands with similar beta values distribution across patients, while computing their mean (a metric that is sensitive to outlier values) we wish to preserve the effect of extreme values. Likewise islands agglomeration, the *K* training-test split and subsequent aggregation of results, aims at alleviating the risk of overfitting when the sample size is small, providing a final weight profile that potentially generalizes better on unseen patients. Finally, NN optimal parametrization is aided by pretraining and network regularization (see Methods). This care for attaining a model with carefully estimated parameters is indeed extremely relevant for the estimation of reliable weight profiles through SHAP. Notably, most of the aforementioned methodological cautions meant to tackle the real-life complexities of epidemiological studies and blood-based DNAm data, can be considered per se valuable algorithmic suggestions whose rationale apply to a broader systems biology research context. For instance, the risk of overfitting resonates with any analysis trying to model complex high-dimensional omic data (s.a., genomic, transcriptomic, metabolomic, gut microbiome, etc.) with ML or DL-based methods when sample size is small. To the best of our knowledge, closely related methods for DNAm data, irrespectively of the specific task, mostly test on large datasets from public biobanks. Similarly, the attentions devoted to the reliability of SHAP-derived explanations, or the peculiar use of the background data to answer precise research question, are relevant aspects that have been largely ignored.

## Conclusion

In conclusion, the contributions of this work are twofold: on the one side, by presenting the results of a case study in the context of BC research, we defend our claims and highlight the need for a paradigm shift from Standard EWAS approach when modeling blood-derived DNAm data in the search for effective TTD biomarkers; on the other, we describe the methodology we applied to effectively achieve the desired effect profile, that has per se several notable and transferrable points of attention. A limit of the present study is the lack of external validation. However, the complex multi-step algorithm based on highly parametrized models we applied here opens the way for future works, that will investigate on what of these algorithms should be transferred as-is on new data (e.g., feature clusters, model parameters, etc.) or retrained from scratch. Nevertheless, the strength of the biological results obtained on the highly challenging case study presented here, and the generalizable methodological cautions, make the present work a relevant milestone in the advancement of blood-based DNAm studies and epidemiological studies in general, whenever robust inference of global effect profiles on a time to event outcome is needed.

## Materials and Methods

### Study sample description

The European Prospective Investigation into Cancer and Nutrition (EPIC) is a large European study on diet and cancer and has been previously described elsewhere [25]. The Italian component of EPIC (EPIC-IT) [26] recruited 47,749 adult volunteers (men and women) at five centres. It is a prospective cohort study with blood samples collected from healthy participants at recruitment. After recruitment, participants were then observed for over 15 years for the insurgence of cancer, cardiovascular diseases and all-cause mortality. At the end of the follow up, all breast, colon and lung cancer cases with available blood sample suitable for epigenetic analyses were paired with an equal number of controls, individually matched on age (±5 years), sex, season of blood collection, center, and length of follow-up.

A detailed description of data collection, DNA methylation measurements and pre-processing and sample filtering can be found in S1 Appendix.

In this work, we focused on the BC case-control study, composed of 248 BC cases and an equal number of matching controls. Additionally, we exploited samples collected for lung cancer (168 control subjects) and colon cancer (140 control subjects) to enlarge our controls group exploited for effects profile weight estimation as described in section Validation through Gene Set Enrichment Analysis. Each subject was described in terms of CpG sites methylation scores, reported as *β*-values, i.e., the proportion of methylated cells per site over the total. Therefore, each CpG site is a continuous value bounded between 0 (no methylation) and 1 (complete methylation). After DNAm data preprocessing, quality control and filtering, we had data for 313,324 CpG sites per subject. In this work we focus on CpG sites corresponding to transcription factor binding sites of EZH2 and SUZ12: two proteins pertaining to the Polycomb Repressive Complex 2 (PRC2). This choice was motivated by previous evidence of the accumulation of DNAm outliers values in these genomic regions, considering also previously described association of the total number of DNAm outliers with BC risk [5]. After filtering on EZH2 and SUZ12 CpG sites, we decided to perform an additional pre-processing by grouping CpG sites into CpG Islands. CpG islands are regions of the genome with a high proportion of CpG dinucleotide repeats in which DNAm is generally conserved. To derive CpG islands *β*-values we aggregated all single sites falling in a specific island by computing their mean. By doing that, we eventually obtained a DNAm representation of 3,807 methylated islands for each subject, that was the input dataset for the algorithm described below.

### Ethics statement

This study was performed according to the principles of the Declaration of Helsinki; all EPIC Italy participants provided written informed consent; the Human Genetics Foundation (HuGeF) Ethics Committee approved the study as reported elsewhere [27].

### Details of the proposed approach

The algorithm we crafted for Deep Survival EWAS on blood DNAm comprises several steps: (i) the Feature Agglomeration of CpG Island methylation profiles, (ii) the non-linear survival modeling of the aggregated features to model the complex relationship between the CpG Islands-derived features and BC TTD and (iii) the estimation of the relevance of each of those features in determining TTD (i.e. their importance in predicting survival risk). A schematic graphical representation of the overall process flow is reported in Fig 1. In this section we will provide a more detailed description of the applied algorithm with theoretical underpinnings when needed.

#### Feature agglomeration

The first step of our approach comprises a hierarchical feature clustering that is spatially independent. This technique is similar to a hierarchical agglomerative clustering procedure, but recursively merges features instead of samples.

Specifically, given the initial input data **X**^(*cases*)^ ∈ ℝ^*N ×Q*^, where *N* is the total number of cases and *Q* the total number of CpG Islands, to identify *J* clusters of CpG Islands we exploited the Euclidean Distance with Ward linkage. Then, we computed the representative value of the j-th agglomerated feature as

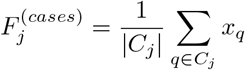

Where *C*_*j*_ is the *j*-th cluster and 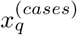 is the *q*-th CpG Island, expressed as *β*-value bounded in [0,1], in the sample of cases. We exploit the same clustering structure (i.e., the same groups *C*_*j*_ of indexes *q*) to compute 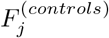 from **X**^(*cases*)^ ∈ ℝ^*N ×Q*^. As mentioned, throughout this Chapter we refer to the aggregated CpG Islands as *features*.

#### Deep Non-Linear Survival Modeling

To model the relationship between the just defined input features and BC TTD we exploit a Deep Survival NN, closely related to DeepSurv [28], to account for the complex non-linear interactions determining the phenotype. In particular, our Deep Survival model is a multi-layer feed-forward NN which predicts the effects of a patient’s covariates on their hazard rate parameterized by the weights of the network *θ*. The input to the network for patient i is its baseline data in terms of DNAm features (*F* ^(*i*)^), while the output 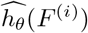 is a single node with a linear activation which estimates its log-risk function. Let *T* be the times to disease and *E* the event indicator, the objective function to optimize becomes:

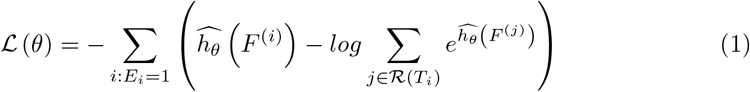

where the risk set ℛ (*t*) = {*i* : *T*_*i*_ ≥ *t*} is the set of patients still at risk of disease at time *t*.

#### Deep Survival Model Architecture and Training

The architecture of the Survival NN was inspired by DeepSurv containing a set of consecutive fully connected layers of decreasing dimensionality (i.e., number of nodes), each followed by a batch normalization layer. The output layer is composed of one node only, that makes a linear combination of the nodes in the second-last layer to predict the log-risk function 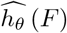. The number of layers and the number of nodes in each layer were optimized as described in Section.

Among other choices to improve Deep Survival Model training and mitigate the risk of suboptimal parametrization, we included an *unsupervised pre-training* step before training our model to minimize the survival loss function. In particular, given a NN model architecture from input to second-last layer (embedding layer) before the single output node, hereby defined encoder, we initialized its parameters through AutoEncoder (AE) Layer-wise pretraining. In this work, we exploit the concept of AEs to identify an initial set of parameters for the Survival NN. To do that, we train progressively deeper AEs: starting from the input dimensionality of the model (input layer), we build an AE with the next layer as bottleneck and we train it to reconstruct the input; then, we retain the parameters from input layer to first layer to build and train the next AE from input to second layer of our Survival NN, and we progress until we have included all available layers in the encoder as bottlenecks.

At the end of this process, the parameters of the encoder are retained as initialization for the Survival NN.

While pretraining is known to aid network regularization, the set of parameters from second-last to output node are initialized at random, therefore we foster their regularization adding a Drop-out layer in between.

#### CpG Islands effects profile estimation through SHAP

To estimate the effect of the CpG Island-derived features on the prediction of the log-risk function we exploit a post-hoc explanation method applied to the deep survival NN. Among the existing methods, SHapley Additive exPlanation (SHAP), by [10], an algorithm that aims at explaining the prediction of an instance by computing the contribution of each feature to the prediction. This contribution is estimated in terms of Shapley regression values, an additive feature attribution method inspired by the coalitional game theory [29]. In the original Shapley formulation, feature impact is defined as the change in the expected value of the model’s output when a feature is observed versus unknown. Given a specific prediction *f* (*x*), we can compute the Shapley values *φ*_*i*_ (*f, x*) using a weighted sum that represents the impact of each feature being added to the model averaged over all possible orders of features being introduced:

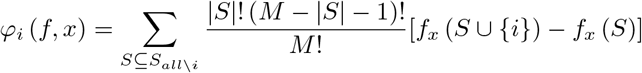

where S is a subset of all the features (*S*_*all*_) used in the model, *M* is the number of features and *f*_*x*_ (*S*) is the prediction for feature values in set S that are marginalized over features that are not included in that set. This computation is prohibitive, therefore SHAP’s authors proposed several sampling-based alternatives to estimate these values. Among them, we chose KernelSHAP [10], that is model agnostic. In KernelSHAP, “Missing features” in a sampled feature coalition are simulated by averaging the model’s prediction over bootstrapped samples with values for these features sampled from the so called ***background* dataset**. This makes the estimation dependent on the choice of such reference, an aspect that we exploited to gather different biological insights by changing the BC cases composition of the background dataset supplied to the method (cf. Section).

In general, in our algorithm, to derive the importance weights that constitute the global effects profile, for each of the *K* splits, we trained the model on the training data and we computed SHAP values of test data w.r.t. the background control group. As SHAP values are computed locally for each observation, we estimate the features’ impact (*w*^(*k*)^) as the mean of the absolute value of local estimates.

Moreover, to reduce computational time of the nested sampling to estimate SHAP values, as suggested by the authors [11], we supplied to the algorithm the background data grouped into 20 centroids, i.e. a sample of 20 representative observations derived from the application of k-means algorithm (with *k* = 20) to the reference sample.

### Performance measures

#### Time-to-event prediction performance

To evaluate the modeling performance of both our Deep Survival NN model and CoxPH we exploited the traditional Harrel’s Concordance Index (CI) [30]. The Harrel CI is a measure of rank correlation between the models’ predicted risk scores and the observed time points. It quantifies how well a model predicts the ordering of patients’ diagnosis times. It can be computed by the following formula:

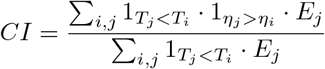

where *η*_*i*_ is the risk score of unit *i*.

#### Weights profiles robustness

To estimate the robustness of the effects profile estimated across K iterations in our Deep Survival EWAS, we exploited the metric developed in [17]. Specifically, for each split k, with *k* = 1, …, *K*, we get a different 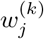 and we rank order features based on these importance scores. To measure weights robustness we measure the dispersion of these K SHAP-derived importance rankings **r**_1_, …, **r**_*K*_. To do that, for each pair of ranked vectors **r**_*i*_ and **r**_*j*_ we compute their dissimilarity as the **Kendall Tau Distance between two rank vectors** [31]. This distance can be expressed as the minimum number of bubble swaps needed to convert one rank vector to the other, divided by the total number of pairs in the vector. Additionally, we impose a 1*/j* penalty on any comparison involving the *j*-th rank as it is more relevant to identify the top predictors of a model, rather than accurately rank all features. Finally, we truncate the rank vectors after the top 10 ranks to reduce computation time. The mean of these 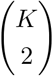 pairwise distances represent the stability of our measurements, that we define Kendall Tau Stability (KT-stability).

### Algorithm Design and Optimization

To optimize the design of the overall pipeline of the algorithm we needed to make several architectural and hyperparameter choices. In particular, we had to optimize jointly the dimensionality of the input to the Survival NN model (i.e. the number of features clusters) and the architecture of the NN itself. To do that, we tested a set of combinations and compared their performances on CI and KT-stability, with the ultimate goal of achieving a reliable set of weights for the effects profile. In particular, we defined a set of input dimensions *J* ∈ [128, 256, 512, 1024], and for each of those dimensions we tested two alternative NN architecture, reducing the number of nodes in half at each subsequent layer down to a final embedding layer of either 16 or 32 nodes. Therefore, in total, we tested 8 NN configurations. The dimensions of NN layers were defined as mentioned in order to aid memory efficiency in model training, and to make these modular architectures more comparable. In total, we tested 8 NN configurations, as reported in S1 Table in Supporting Information. For pre-training, model parameters were initialized randomly following the approach in Glorot et al. The MSE loss function was optimized by mini-batch stochastic gradient descent with momentum with a learning rate of 0.1 and a momentum parameter of 0.9, in accordance with the most common hyperparameters definition for this optimizer. The mini-batch size was set to 32, counting for around 15% of the training set. We trained the model for 150 epochs. After pre-training, the resulting parameters were exploited as initialization parameters for the overall survival NN, fine-tuned through gradient descent of the previously defined survival loss function, while the last layer was initialized randomly. The optimization was performed with adaptive moment estimation, inspired by DeepSurv, with a learning rate of 0.0001, a mini-batch size of 32 samples and a drop-out probability of the last fully connected layer of 0.1. We fine-tuned the whole network for 150 epochs leaving all other hyperparameters to default. To identify the best combination of input dimensionality and survival model, we first defined the 4 grouped input datasets by varying J in feature agglomeration, then we run the entire pipeline from NN pretraining to SHAP weights estimation (using all BC controls as background data) repeated on 10 random data splits, comparing the average results (cf. S1 Table). As an additional decision support resource, we compared the distributions of CpG Islands grouped in each aggregated feature for the different K values (S4 Fig). As none of the alternative showed significantly superior performances on the two metrics together, we opted for the smaller model (i.e. 128 nodes in the input and 16 nodes in the last fully connected layer before output) that granted comparable results to the others with significantly less parameters to train. Moreover, the size distribution with input granularity of 128 features (cf. S4 Fig) shows an average of features’ size between 19 and 35 CpG Islands per feature, which seemed a balanced and reasonable clustering considering the input of almost 4,000 Islands.

## Supporting information

**S1 Appendix. Detailed Data Description**. Supporting information on cohort description and DNA methylation data extraction and preprocessing.

**S1 Table. Optimization Results** Performance in terms of Kendall-Tau Stability (robustness) and Harrell C-Index (survival prediction performance) for all the Deep Survival Network architecture trained in optimization. Performance is averaged across K splits; 95% confidence intervals are reported in parentheses. In the first column (Input) the input shape (*J*), referring to the granularity of the CpG Island agglomeration. In the second column (Latent) the dimensionality of the Survival NN layer before output. In the second column (Architecture), the list of layers with respective number of nodes, up until before output layer (one single output node for all architectures).

**S2 Table. Deep Survival EWAS Effects profile estimation w.r.t. reference groups**. Estimated CpG islands effects profiles. Each sheet in the file contains the results for one reference group (respectively: (1) BC controls, (2) Matching Controls, (3) All controls, (4) All controls with cases). The first column reports the CpG Island name, the second reports the Feature they are agglomerated into. Third column reports the effect weight (*w*), while last column reports the ranking of the feature in terms of importance (i.e. ordered by descending magnitude of effect weight associated to the feature).

**S1 File. Kaplan-Meyer Plots of** *F*_120_ **CpG Islands**.

**S3 Table. Deep Survival EWAS Gene Set Enrichment Analyses Results**. Deep Survival EWAS GSEA results for all four reference groups. Each sheet in the file contains the results for one reference group (respectively: (a) BC controls, (b) Matching Controls, (c) All controls, (d) All controls with cases). The first column reports the KEGG pathway, followed by pathway code and empirical p-value (last column).

**S4 Table. Standard EWAS association results**. Results of independent modeling of CpG Islands w.r.t. TTD. The table reports p-values and test statistics obtained by modeling one univariate CoxPH for each island.

**S5 Table. Standard EWAS Gene Set Enrichment Analysis results**. Results of wighted GSEA for Standard EWAS approach, where weights were determined by the test statistics in the univariate CoxPH performed independently for each CpG Island. The first column reports the KEGG pathway, followed by pathway code and empirical p-value (last column).

**S1 Fig. Standard EWAS weights profile**. Weights profile for Standard EWAS approach, where each CpG Island is associated with the p-value of the test statistic in an independent CoxPH model. Panel A reports the p-values without Bonferroni adjustment, panel B reports the same p-values after Bonferroni adjustment. The red line denotes the p-value threshold of 0.05.

**S6 Table. Multivariate Cox Proportional Hazards results**. Performance in terms of Kendall-Tau Stability (robustness) and Harrell C-Index (survival prediction performance) for all Multivariate CoxPH models with different input dimensions. Performance is averaged across K splits; 95% confidence intervals are reported in parentheses. In the first column (Input) the input shape (*J*), referring to the granularity of the CpG Island agglomeration.

**S2 Fig. Predictive Performance comparison**. Predictive Performance (Harrel CI) of all the tested Deep Survival NN architectures (blue) and all multivariate CoxPH models fitted. The values in parenthesis for Deep Survival NNs represent the number of input nodes (i.e. the granularity of features’ clusters) and the number of nodes in the last layer before the output. Whereas the parenthesis for CoxPH models report the granularity of the input features’ clusters. Dots represent the average performance value, while bands report the confidence intervals around the mean computed on the K=10 splits.

**S3 Fig. Weights stability comparison**. Importance weights stability performance (KT-stability) of all the tested Deep Survival NN architectures (blue) and all multivariate CoxPH models fitted. The values in parenthesis for Deep Survival NNs represent the number of input nodes (i.e. the granularity of features’ clusters) and the number of nodes in the last layer before the output. Whereas the parenthesis for CoxPH models report the granularity of the input features’ clusters. Dots represent the average performance value, while bands report the confidence intervals around the mean computed on the K=10 splits.

**S4 Fig. Features’ dimension distributions for varying** *J*. Distribution of dimensions of the features for each input granularity *J* (i.e. 128, 256, 512, 1024), in terms of number of CpG Island they group.

## Acknowledgments

We thank the EPIC Italy research group (Carlotta Sacerdote, Vittorio Krogh, Domenico Palli, Salvatore Panico, Rosario Tumino, Paolo Vineis and their collaborators) for giving us access to the data of the BC study nested in the cohort.

## Data Availability

DNAm data used in this study are available at the GEO repository under the accession number GSE51032.

## References

1. Yang X, Yan L, Davidson NE. DNA methylation in breast cancer. Endocrine-related cancer. 2001;8(2):115–127.

2. Das PM, Singal R. DNA methylation and cancer. Journal of clinical oncology. 2004;22(22):4632–4642.

3. Bodelon C, Ambatipudi S, Dugué PA, Johansson A, Sampson JN, Hicks B, et al. Blood DNA methylation and breast cancer risk: a meta-analysis of four prospective cohort studies. Breast Cancer Research. 2019;21(1):1–9.

4. Hüls A, Czamara D. Methodological challenges in constructing DNA methylation risk scores. Epigenetics. 2020;15(1-2):1–11.

5. Gagliardi A, Dugué PA, Nøst TH, Southey MC, Buchanan DD, Schmidt DF, et al. Stochastic epigenetic mutations are associated with risk of breast cancer, lung cancer, and mature b-cell neoplasms. Cancer Epidemiology and Prevention Biomarkers. 2020;29(10):2026–2037.

6. Levy JJ, Titus AJ, Petersen CL, Chen Y, Salas LA, Christensen BC. MethylNet: an automated and modular deep learning approach for DNA methylation analysis. BMC bioinformatics. 2020;21(1):1–15.

7. Zheng C, Xu R. Predicting cancer origins with a DNA methylation-based deep neural network model. PloS one. 2020;15(5):e0226461.

8. Macías-García L, Martínez-Ballesteros M, Luna-Romera JM, García-Heredia JM, García-Gutiérrez J, Riquelme-Santos JC. Autoencoded DNA methylation data to predict breast cancer recurrence: Machine learning models and gene-weight significance. Artificial Intelligence in Medicine. 2020;110:101976.

9. Liu B, Liu Y, Pan X, Li M, Yang S, Li SC. DNA methylation markers for pan-cancer prediction by deep learning. Genes. 2019;10(10):778.

10. Lundberg SM, Lee SI. A unified approach to interpreting model predictions. In: Proceedings of the 31st international conference on neural information processing systems; 2017. p. 4768–4777.

11. Lundberg SM, Erion G, Chen H, DeGrave A, Prutkin JM, Nair B, et al. From local explanations to global understanding with explainable AI for trees. Nature machine intelligence. 2020;2(1):56–67.

12. Liu H, Wu X, Zhang S. Feature selection using hierarchical feature clustering. In: Proceedings of the 20th ACM international conference on Information and knowledge management; 2011. p. 979–984.

13. Yousefi-Azar M, Varadharajan V, Hamey L, Tupakula U. Autoencoder-based feature learning for cyber security applications. In: 2017 International joint conference on neural networks (IJCNN). IEEE; 2017. p. 3854–3861.

14. Levy JJ, Chen Y, Azizgolshani N, Petersen CL, Titus AJ, Moen EL, et al. MethylSPWNet and MethylCapsNet: Biologically Motivated Organization of DNAm Neural Network, Inspired by Capsule Networks. bioRxiv. 2021; p. 2020–08.

15. Mallik S, Seth S, Bhadra T, Zhao Z. A linear regression and deep learning approach for detecting reliable genetic alterations in cancer using dna methylation and gene expression data. Genes. 2020;11(8):931.

16. Kent WJ, Sugnet CW, Furey TS, Roskin KM, Pringle TH, Zahler AM, et al. The human genome browser at UCSC. Genome research. 2002;12(6):996–1006.

17. Liu B, Udell M. Impact of Accuracy on Model Interpretations. arXiv preprint 201109903. 2020;.

18. Charmpi K, Ycart B. Weighted Kolmogorov Smirnov testing: an alternative for gene set enrichment analysis. Statistical applications in genetics and molecular biology. 2015;14(3):279–293.

19. Khodabandehlou N, Mostafaei S, Etemadi A, Ghasemi A, Payandeh M, Hadifar S, et al. Human papilloma virus and breast cancer: the role of inflammation and viral expressed proteins. BMC cancer. 2019;19(1):1–11.

20. Su J, Yan D, Wu S, et al. Epstein-Barr virus infection and increased sporadic breast carcinoma risk: a meta-analysis. Medical Principles and Practice. 2020;29(2):195–200.

21. Ortega MA, Fraile-Martínez O, Asúnsolo Á, Buján J, García-Honduvilla N, Coca S. Signal transduction pathways in breast cancer: the important role of PI3K/Akt/mTOR. Journal of oncology. 2020;2020.

22. Azimi I, Roberts-Thomson S, Monteith G. Calcium influx pathways in breast cancer: opportunities for pharmacological intervention. British journal of pharmacology. 2014;171(4):945–960.

23. Xu Z, Sandler DP, Taylor JA. Blood DNA methylation and breast cancer: a prospective case-cohort analysis in the sister study. JNCI: Journal of the National Cancer Institute. 2020;112(1):87–94.

24. Liu Y, Li X, Aryee MJ, Ekström TJ, Padyukov L, Klareskog L, et al. GeMes, Clusters of DNA Methylation under Genetic Control, Can Inform Genetic and Epigenetic Analysis of Disease. The American Journal of Human Genetics. 2014;94(4):485–495. doi:https://doi.org/10.1016/j.ajhg.2014.02.011.

25. Gonzalez CA. The European prospective investigation into cancer and nutrition (EPIC). Public health nutrition. 2006;9(1a):124–126.

26. Palli D, Berrino F, Vineis P, Tumino R, Panico S, Masala G, et al. A molecular epidemiology project on diet and cancer: the EPIC-Italy Prospective Study. Design and baseline characteristics of participants. Tumori Journal. 2003;89(6):586–593.

27. Fiorito G, McCrory C, Robinson O, Carmeli C, Rosales CO, Zhang Y, et al. Socioeconomic position, lifestyle habits and biomarkers of epigenetic aging: a multi-cohort analysis. Aging (Albany NY). 2019;11(7):2045.

28. Katzman JL, Shaham U, Cloninger A, Bates J, Jiang T, Kluger Y. DeepSurv: personalized treatment recommender system using a Cox proportional hazards deep neural network. BMC medical research methodology. 2018;18(1):1–12.

29. Shapley LS. A value for n-person games. In: Contributions to the Theory of Games. vol. 2. Princeton University Press; 1953. p. 307–317.

30. Harrell FE, Califf RM, Pryor DB, Lee KL, Rosati RA. Evaluating the yield of medical tests. Jama. 1982;247(18):2543–2546.

31. Kumar R, Vassilvitskii S. Generalized distances between rankings. In: Proceedings of the 19th international conference on World wide web; 2010. p. 571–580.

