## Supplementary material for "A Deep Survival EWAS approach estimating risk profile based on pre-diagnostic DNA methylation: an application to Breast Cancer time to diagnosis": S1 Appendix

### 1. DETAILED COHORT DESCRIPTION

The European Prospective Investigation into Cancer and Nutrition (EPIC) is a large European study on diet and cancer and has been previously described elsewhere [1]. The Italian component of EPIC (EPIC-IT) [2] recruited 47,749 adult volunteers (men and women) at five centres: Varese and Turin in northern Italy, Florence in central Italy and Naples and Ragusa in southern Italy. All participants signed an informed consent form and completed two questionnaires: one about dietary habit (food-frequency) and one about lifestyle, with information on education, socioeconomic status, occupation, history of previous illnesses and surgery, lifetime tobacco use and alcohol consumption and physical activity. EPIC-IT database records were linked to cancer and regional mortality registries after database quality control. All EPIC-IT centres except Naples are covered by population-based cancer registries. In Naples, follow-up information was collected from electronic hospital discharge records and also by periodic personal contact with participants. The study was approved by the ethical review boards of the International Agency for Research on Cancer, and of the collaborating institutions responsible for subject recruitment in each of the EPIC recruitment centres. All centres for EPIC Italy, except for Florence are represented in LIFEPATH.

#### DNA methylation measurements and pre-processing

DNA from blood samples were extracted from buffy coats using the QIAsymphony DNA Midi Kit (Qiagen, Crawley, UK). Bisulphite conversion of 500 ng of each sample was performed using the EZ-96 DNA Methylation-Gold™ Kit according to the manufacturer's protocol (Zymo Research, Orange, CA). Then, bisulfite-converted DNA was used for hybridization on the Infinium HumanMethylation 450 BeadChip, following the Illumina Infinium HD Methylation protocol. Briefly, a whole genome amplification step was followed by enzymatic end-point fragmentation and hybridization to HumanMethylation 450 BeadChips at 48°C for 17 h, followed by single nucleotide extension. The incorporated nucleotides were labeled with biotin (ddCTP and ddGTP) and 2,4-dinitrophenol (DNP) (ddATP and ddTTP). After the extension step and staining, the BeadChip was washed and scanned using the Illumina HiScan SQ scanner. The intensities of the images were extracted using the GenomeStudio (v.2011.1) Methylation module (1.9.0) software, which normalizes within-sample data using different internal controls that are present on the HumanMethylation 450 BeadChip and internal background probes. The methylation score for each CpG was represented as a $\beta$-value according to the fluorescent intensity ratio representing any value between 0 (unmethylated) and 1 (completely methylated).

We extracted raw fluorescence intensities data from 'idat' files and performed background subtraction, colour bias correction, and fluorescence intensities normalization using in house developed R script described elsewhere. Samples were excluded if the bisulphite conversion fluorescence intensity was low (i.e. less than 10,000 for both type-I and type-II probes). Methylation measures were set to missing if the detection p-value was higher than 0.01. Finally, samples and CpG sites with low call rate (<95%) were removed prior to the analysis, as long with probes with non-unimodal distribution (probably methQTL sites). Probe design bias was corrected using the beta-mixture quantile (BMIQ) normalization procedure implemented in the 'wateRmelon' R package [3]. Known batch effect by plate and position on the Illumina Beadchip was removed before statistical analyses using the ComBat algorithm described by Johnson et al. (ComBat function in the "sva" R package) [4]. Finally, the leukocyte composition in each sample was estimated according to Houseman's method [5].

The sample with measured DNA methlyation included individuals from three nested case-control studies on breast, colon and lung cancer. Participants were sampled from the 47,749 participants of the EPIC Italy cohort and included 354 incident breast cancer cases, 169 incident colon cancer cases and 192 incident lung cancer cases and their matched controls. Controls were individually matched on age ($\pm$5 years), sex, season of blood collection, center, and length of follow-up. We excluded cases whose diagnosis was made less than one year since blood draw. Overall, after DNA methylation data quality controls and sample filtering, and excluding samples from Florence Center, for this study we were left with 248 breast cancer cases, 168 lung cancer cases and 140 colon cancer cases, with all the respective matching controls.

### REFERENCES

1. C. A. Gonzalez, "The european prospective investigation into cancer and nutrition (epic)," Public health nutrition **9**, 124–126 (2006).
2. D. Palli, F. Berrino, P. Vineis, R. Tumino, S. Panico, G. Masala, C. Saieva, S. Salvini, M. Cerati, V. Pala *et al.*, "A molecular epidemiology project on diet and cancer: the epic-italy prospective study. design and baseline characteristics of participants," Tumori J. **89**, 586–593 (2003).
3. R. Pidsley, C. C. Y Wong, M. Volta, K. Lunnon, J. Mill, and L. C. Schalkwyk, "A data-driven approach to preprocessing illumina 450k methylation array data," BMC genomics **14**, 1–10 (2013).
4. W. E. Johnson, C. Li, and A. Rabinovic, "Adjusting batch effects in microarray expression data using empirical bayes methods," Biostatistics **8**, 118–127 (2007).
5. W. P. Accomando, J. K. Wiencke, E. A. Houseman, H. H. Nelson, and K. T. Kelsey, "Quantitative reconstruction of leukocyte subsets using dna methylation," Genome biology **15**, 1–12 (2014).
