## Supplementary figures and images for "A Deep Survival EWAS approach estimating risk profile based on pre-diagnostic DNA methylation: an application to Breast Cancer time to diagnosis"

### S1 Fig.

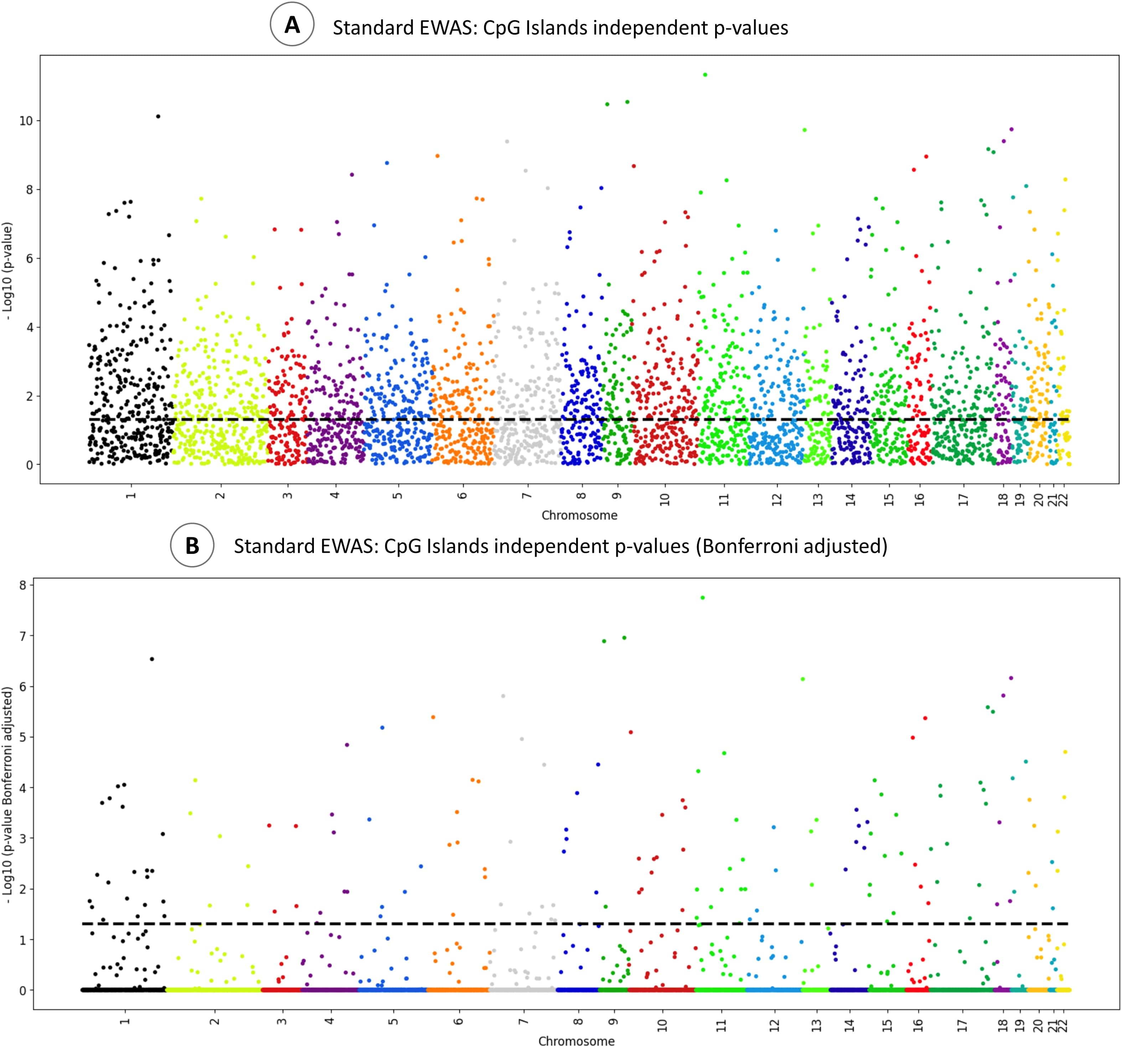

### S1 File

#### Kaplan-Meyer Plots of CpG Islands in $F_{120}$

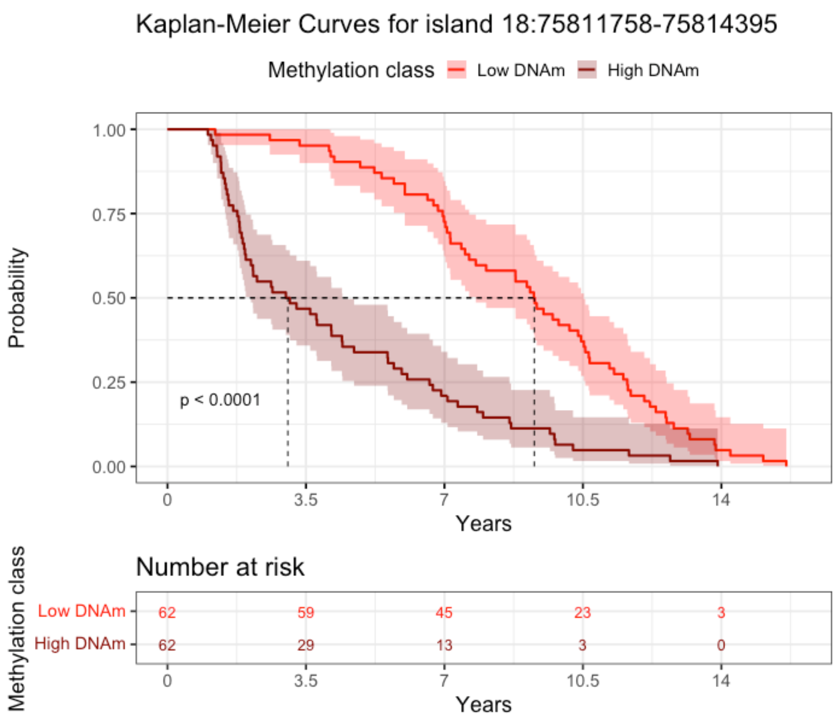


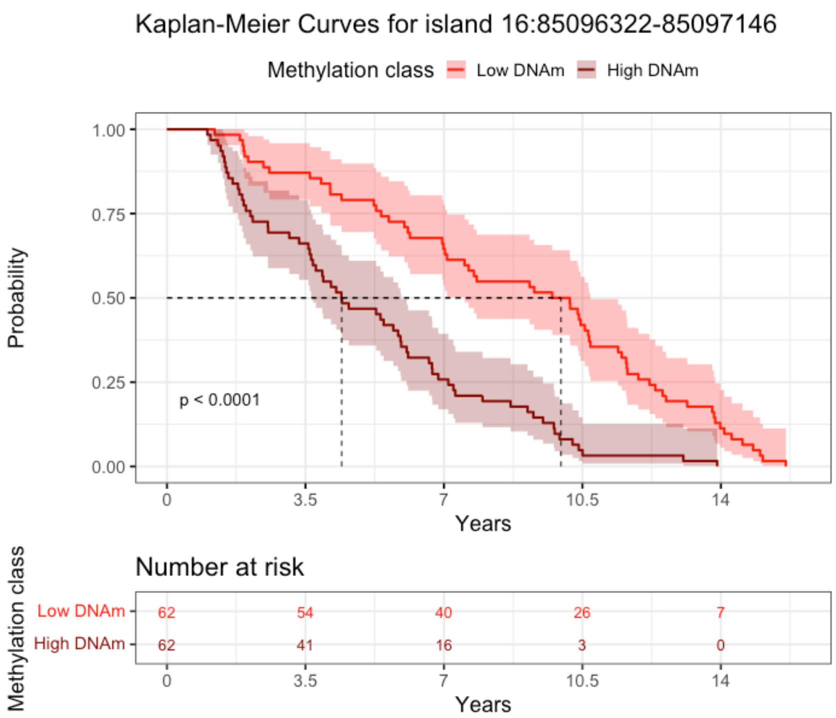


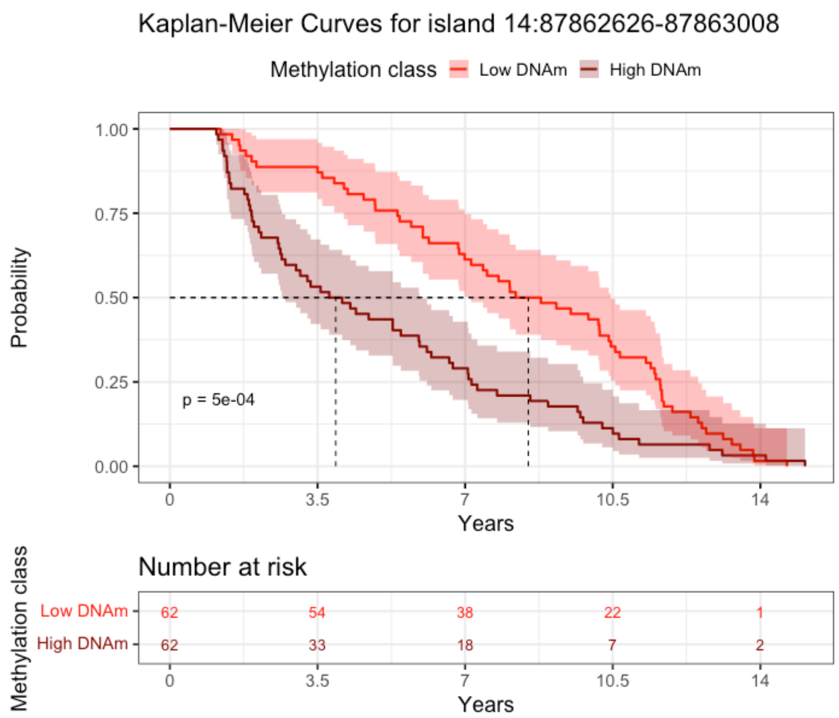


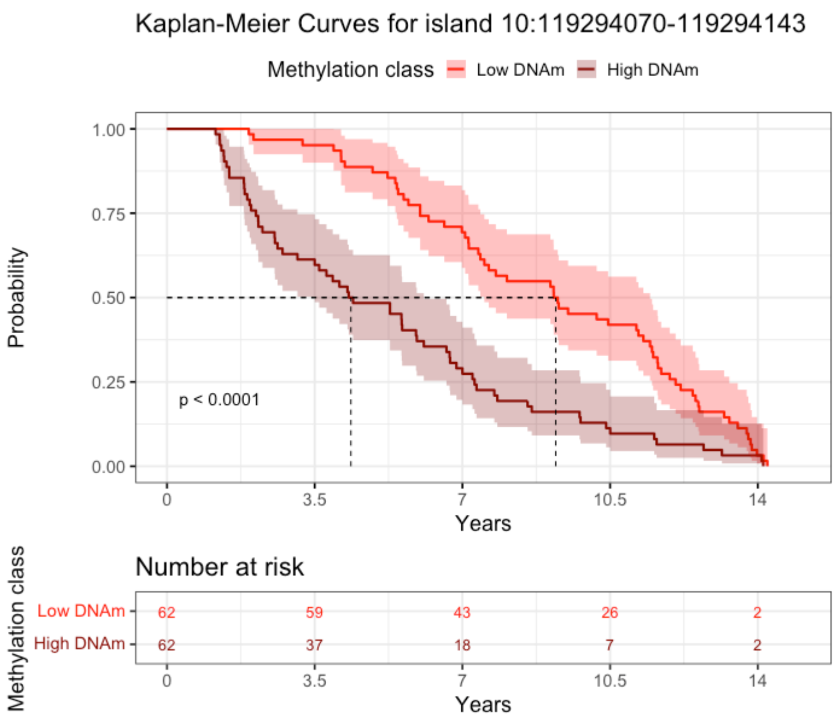


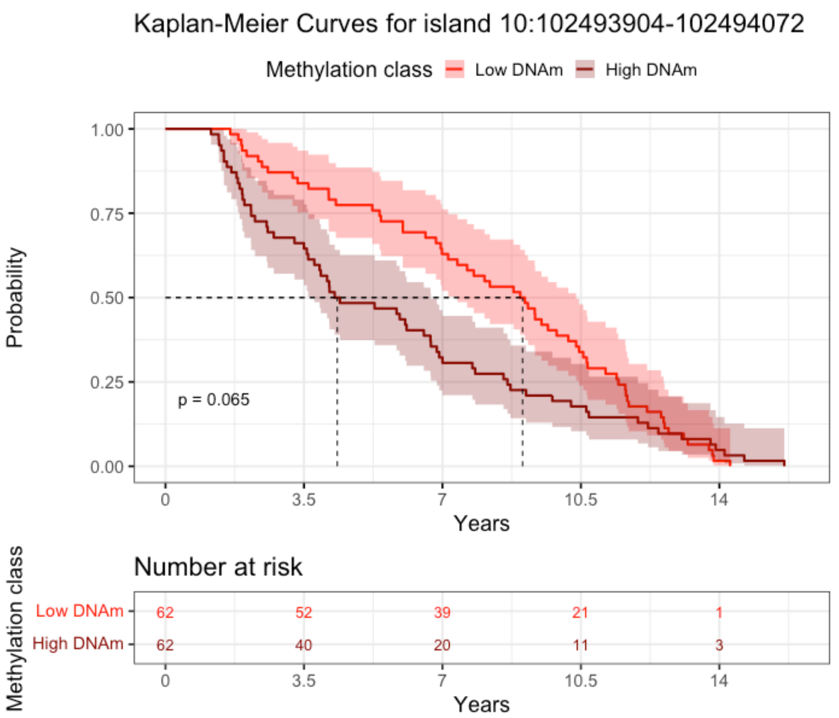


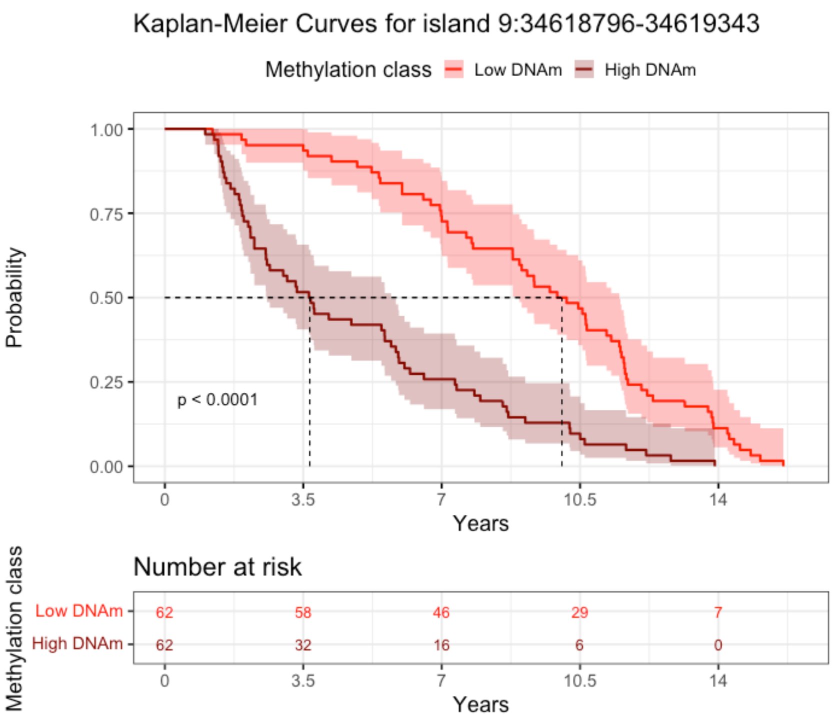

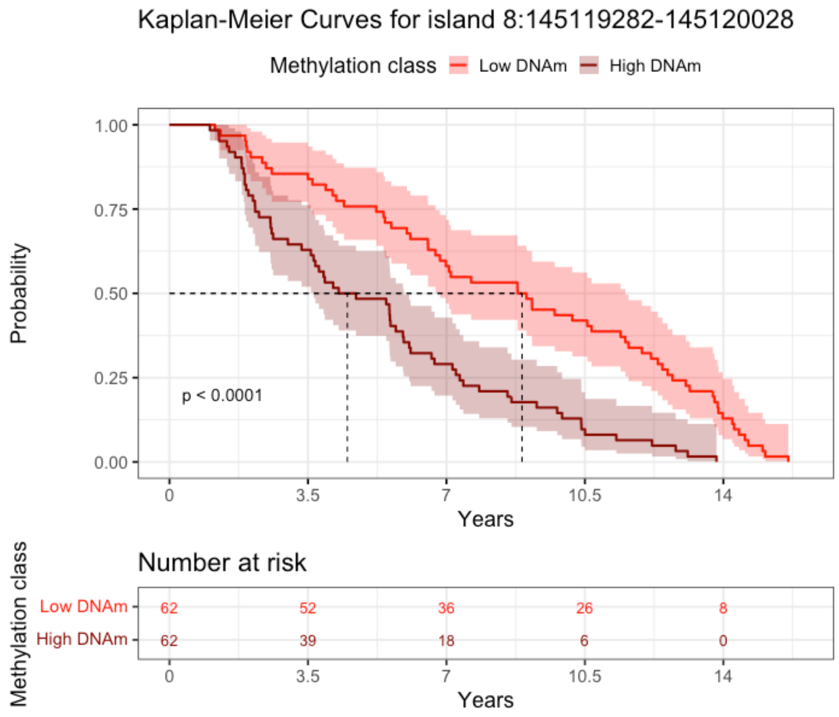

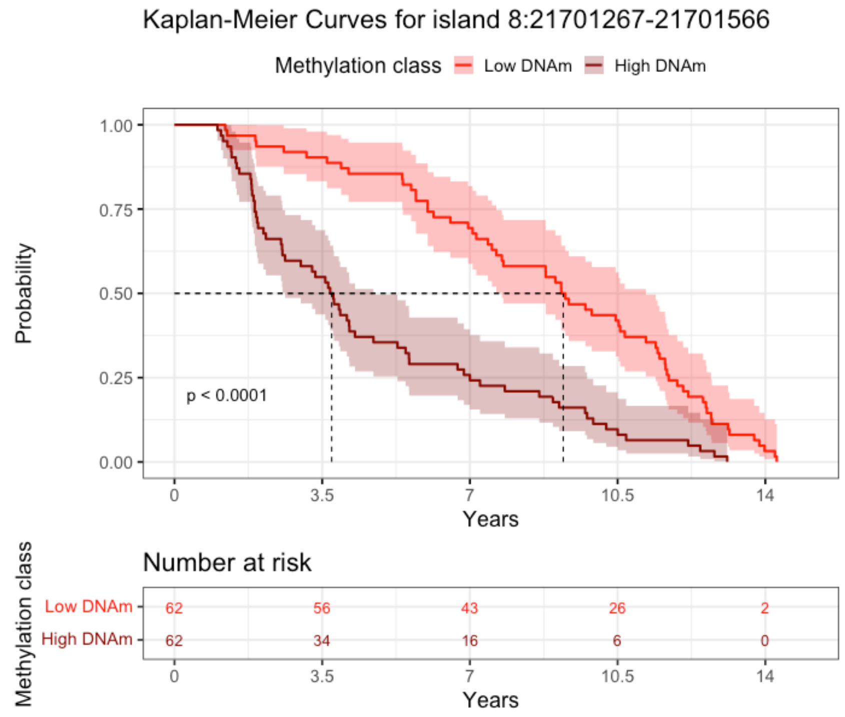

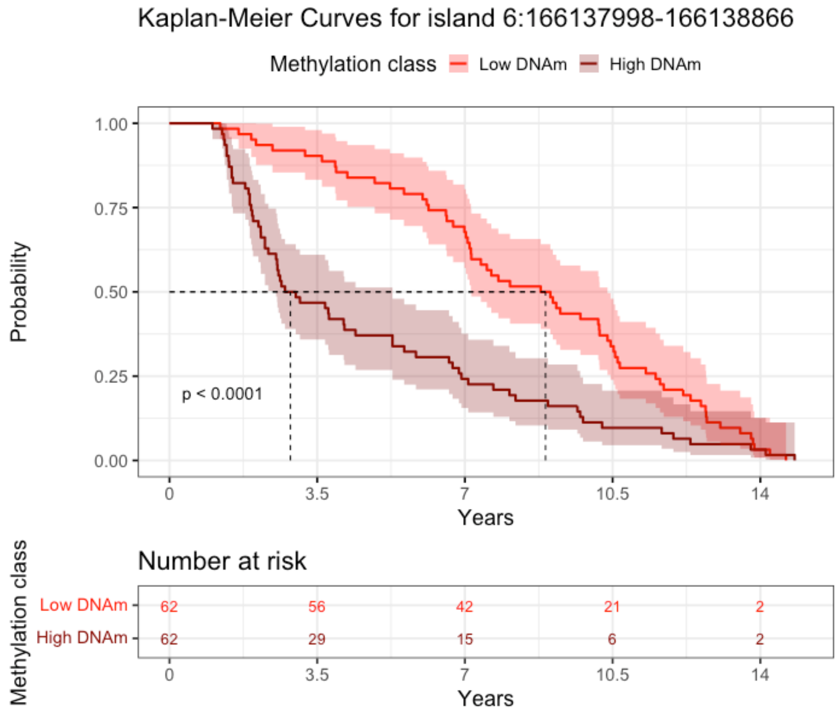

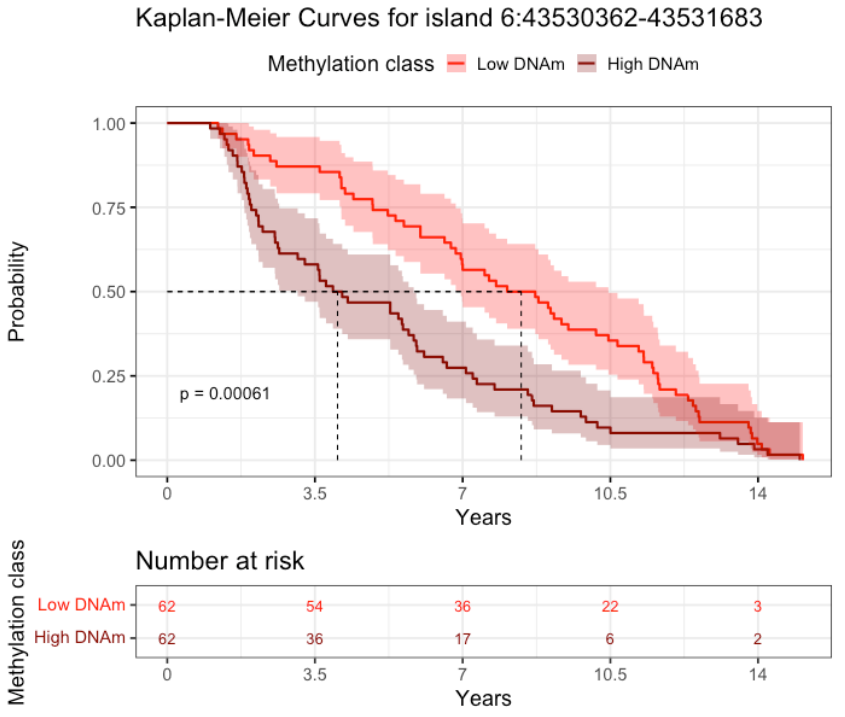

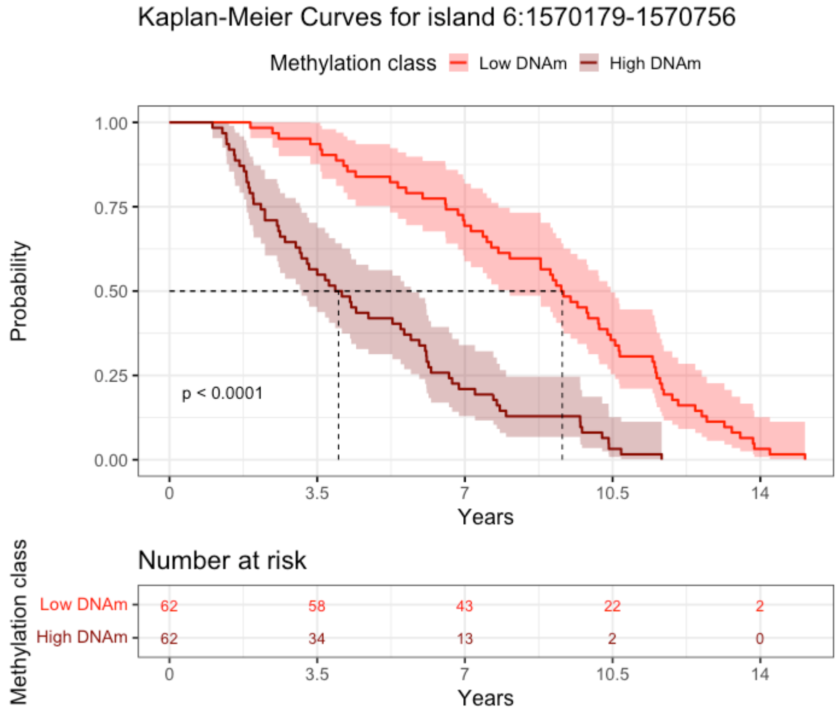

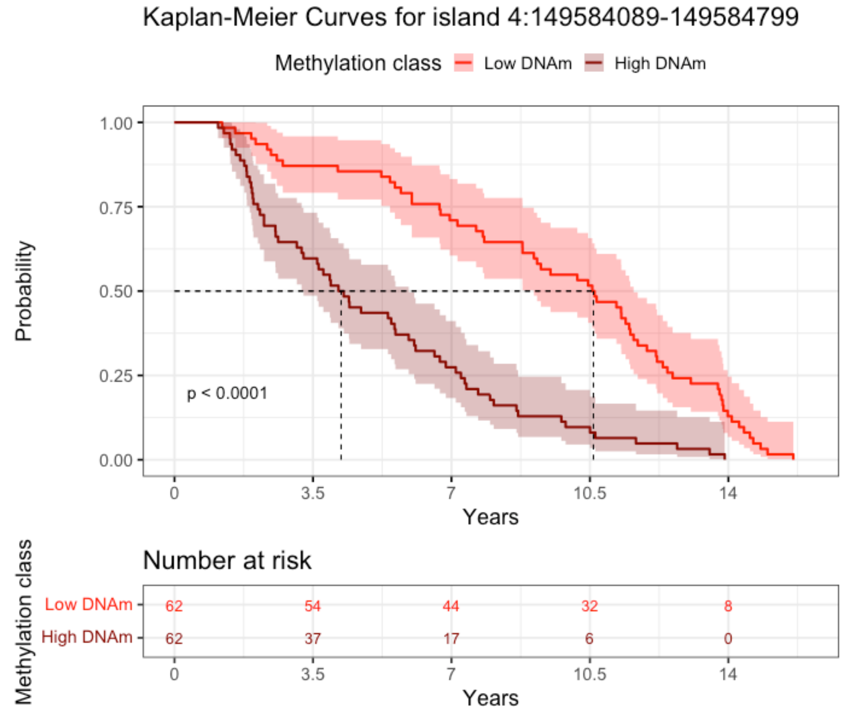

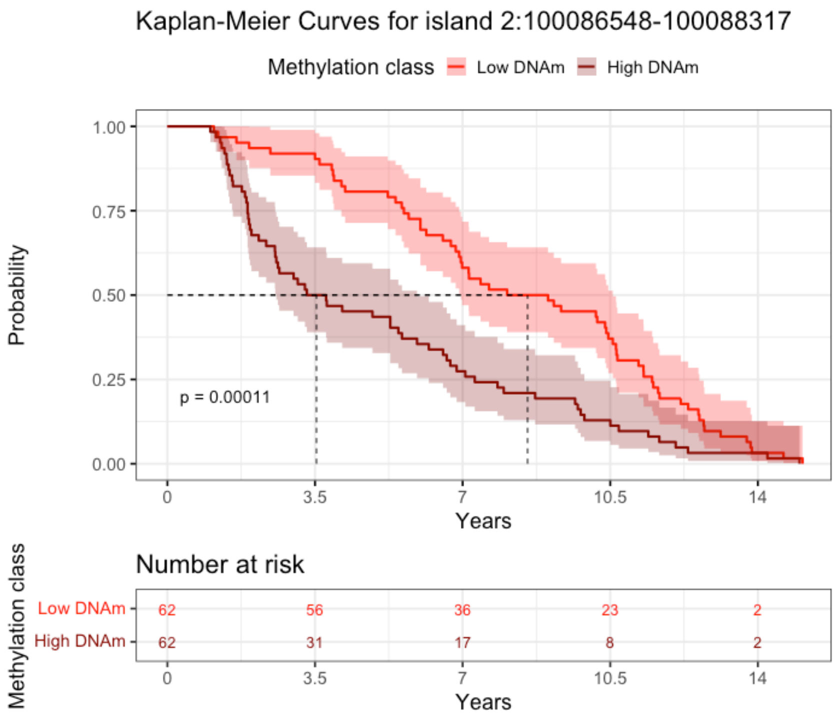

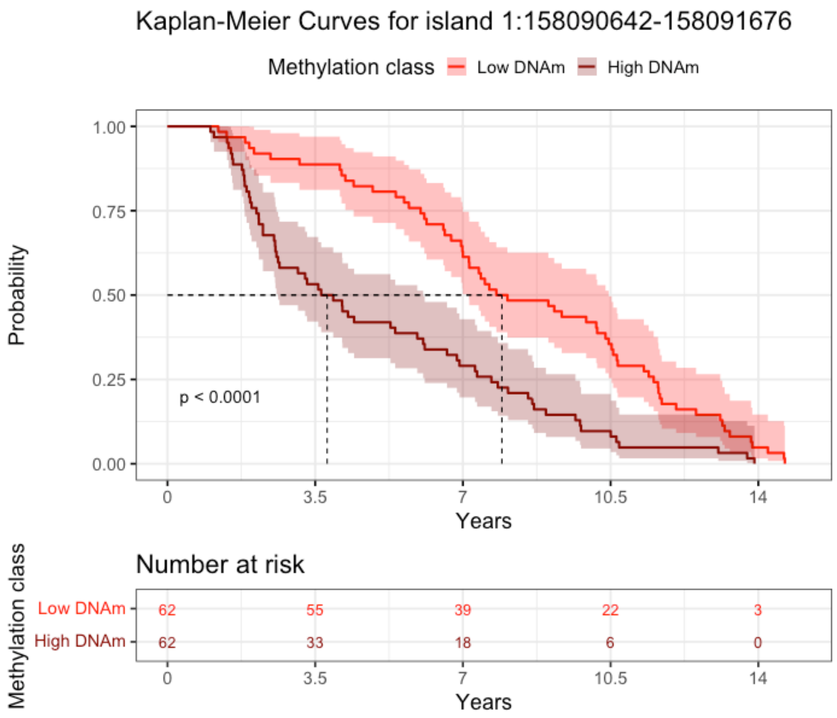

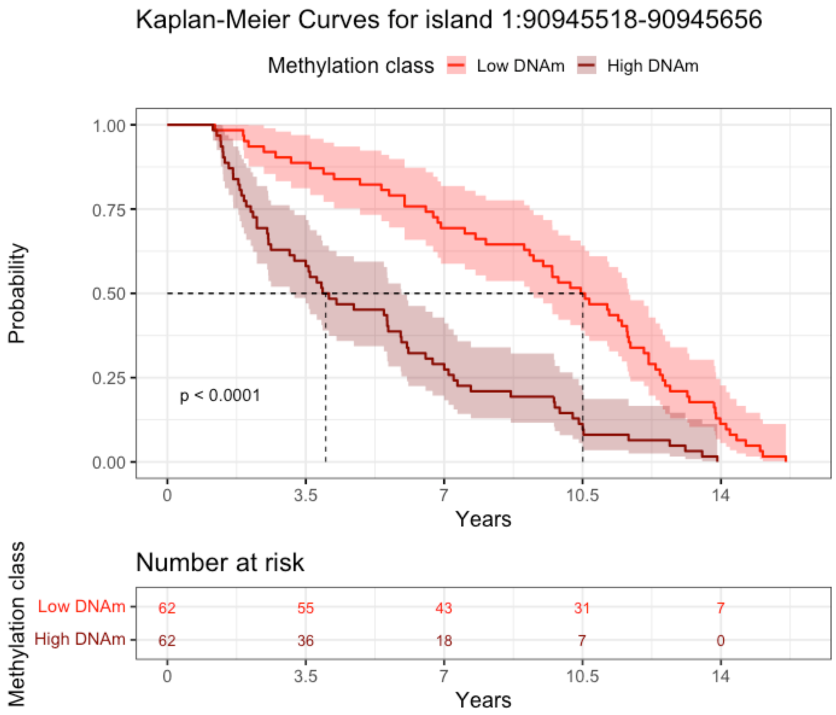

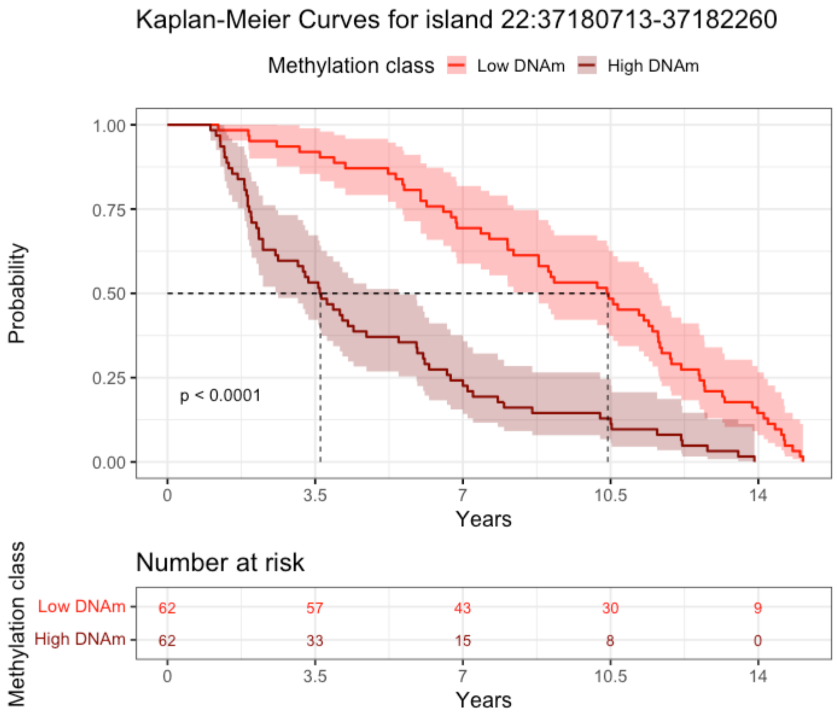

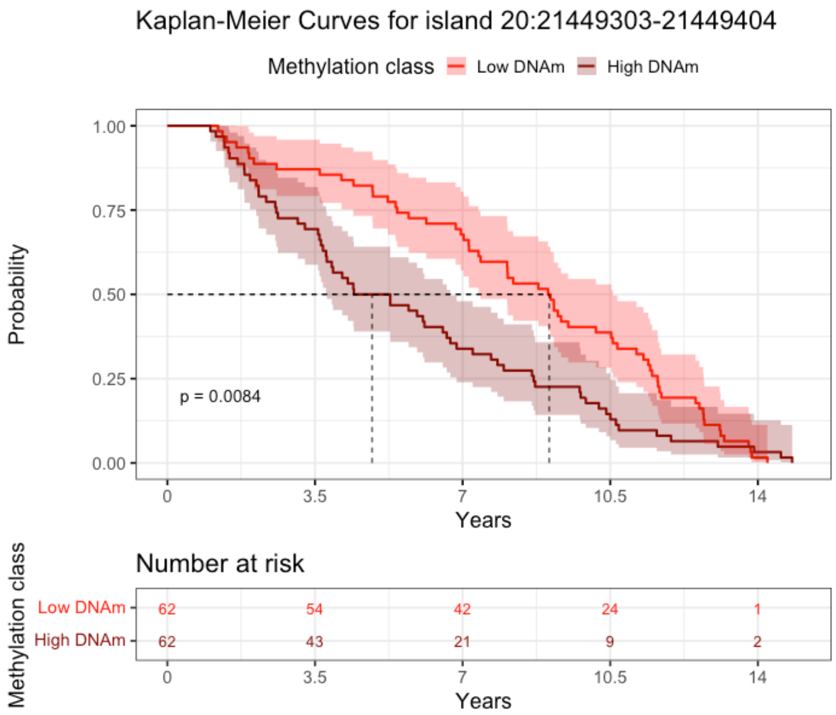

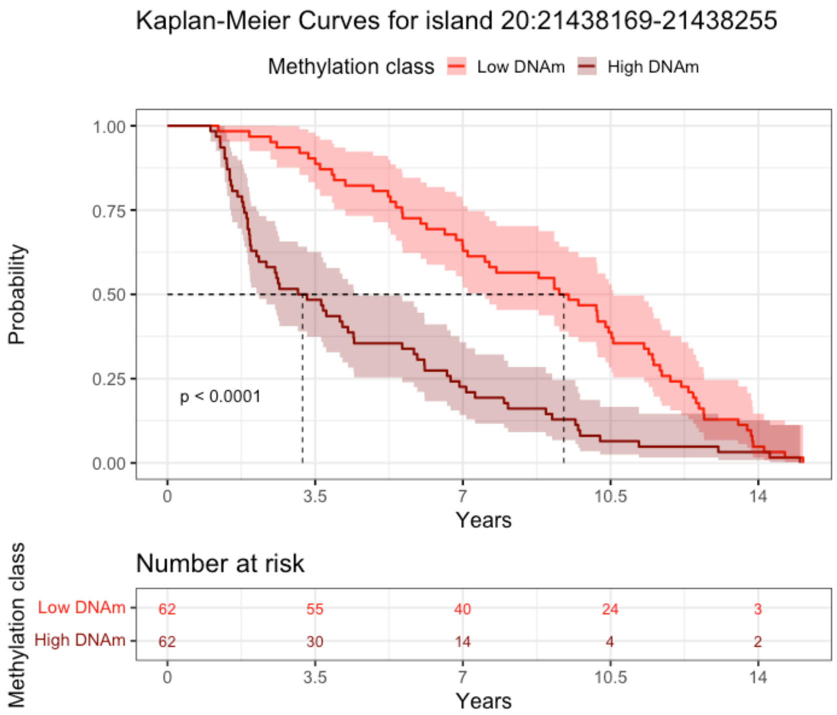

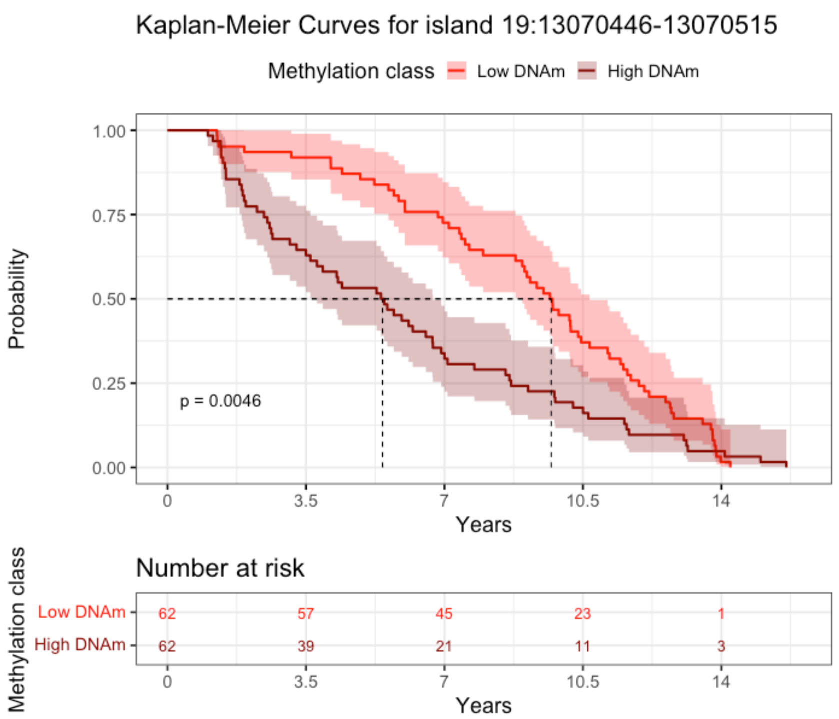

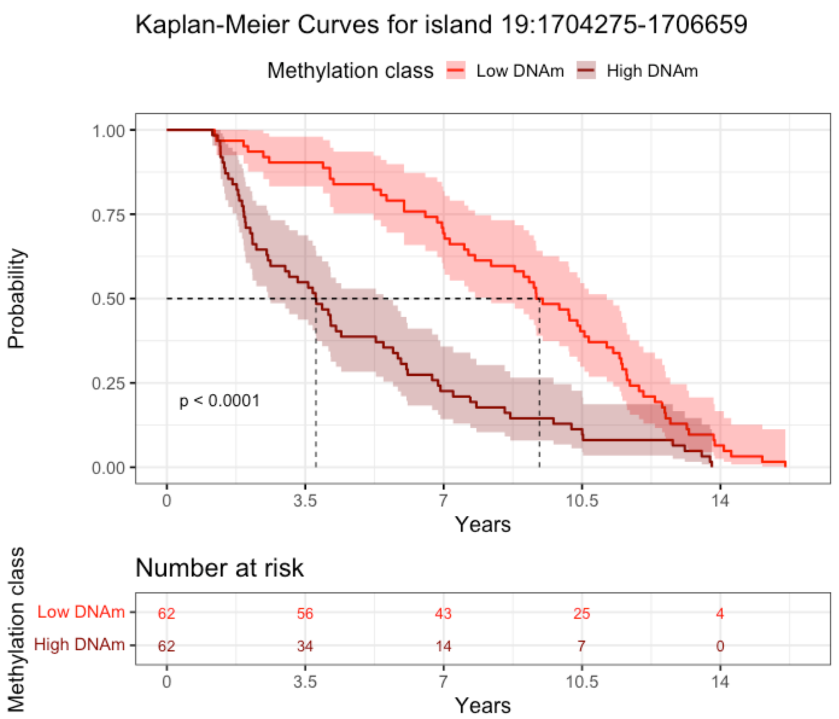

### S2 Fig.

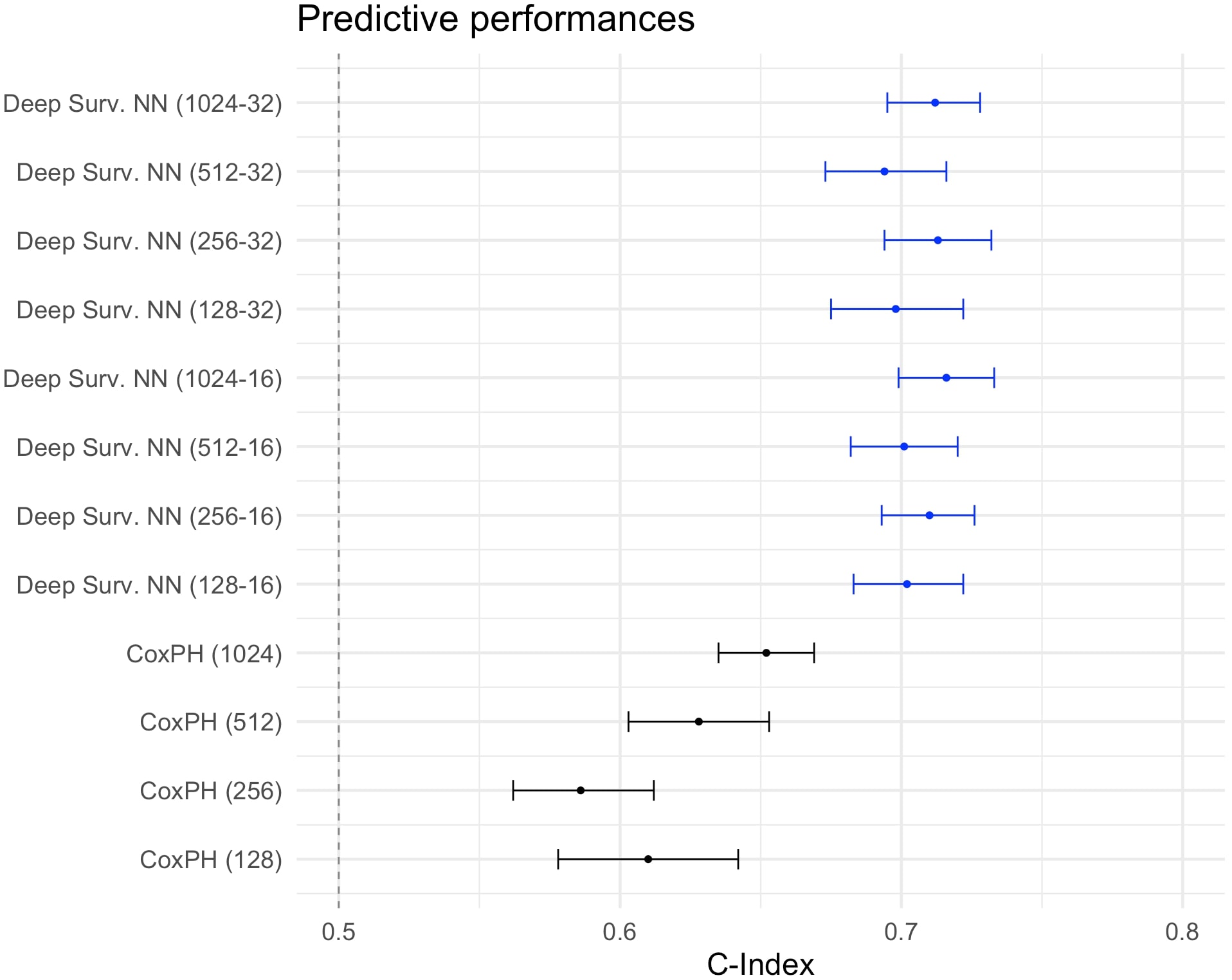

### S3 Fig.

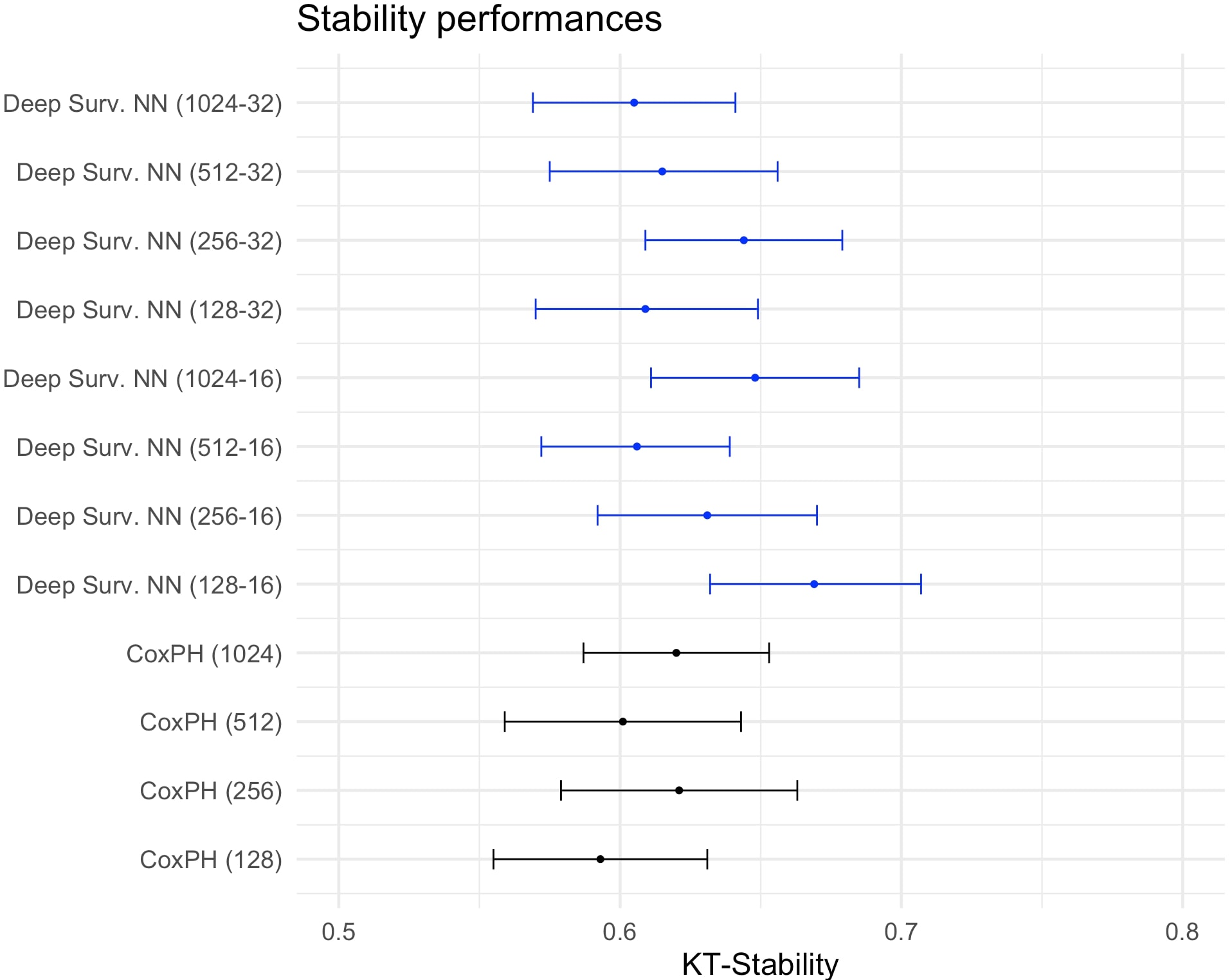

### S4 Fig.

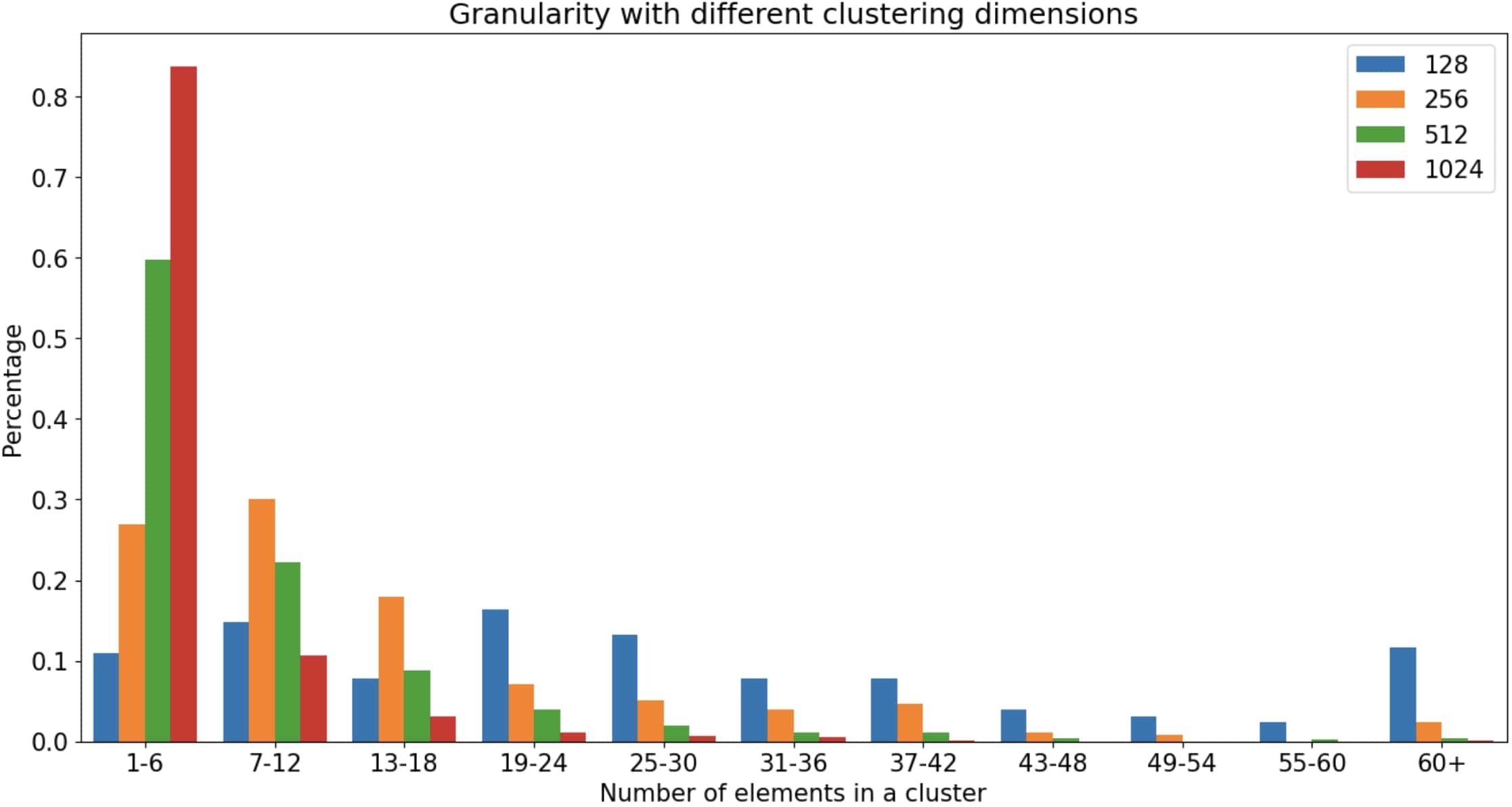
