## Supplementary material for "A Deep Survival EWAS approach estimating risk profile based on pre-diagnostic DNA methylation: an application to Breast Cancer time to diagnosis": S1 Table

**Table S1**

| Input | Architecture (nodes by layer) | KT Stab. (+-95% interval) | C-Index (+-95% interval) |
|---|---|---|---|
| 128 | 128-64-32-16 | 0.669 (+- 0.036) | 0.702 (+- 0.019) |
| 128 | 128-64-32 | 0.609 (+- 0.038) | 0.698 (+- 0.023) |
| 256 | 256-128-64-32-16 | 0.631 (+- 0.039) | 0.710 (+- 0.016) |
| 256 | 256-128-64-32 | 0.644 (+- 0.035) | 0.713 (+- 0.019) |
| 512 | 512-256-128-64-32-16 | 0.606 (+- 0.033) | 0.701 (+- 0.019) |
| 512 | 512-256-128-64-32 | 0.615 (+- 0.040) | 0.694 (+- 0.021) |
| 1024 | 1024-512-256-128-64-32-16 | 0.648 (+- 0.037) | 0.716 (+- 0.017) |
| 1024 | 1024-512-256-128-64-32 | 0.605 (+- 0.036) | 0.712 (+- 0.016) |
