## Supplementary material for "A Deep Survival EWAS approach estimating risk profile based on pre-diagnostic DNA methylation: an application to Breast Cancer time to diagnosis": S6 Table

Table S6

| Input | KT Stab. (+-95% interval) | C-Index (+-95% interval) |
|---|---|---|
| 128 | 0.593 (+- 0.038) | 0.610 (+- 0.032) |
| 256 | 0.621 (+- 0.042) | 0.586 (+- 0.024) |
| 512 | 0.601 (+- 0.042) | 0.628 (+- 0.025) |
| 1024 | 0.620 (+- 0.033) | 0.652 (+- 0.017) |
